# BANDITS: Bayesian differential splicing accounting for sample-to-sample variability and mapping uncertainty

**DOI:** 10.1101/750018

**Authors:** Simone Tiberi, Mark D Robinson

## Abstract

Alternative splicing is a biological process during gene expression that allows a single gene to code for multiple proteins. However, splicing patterns can be altered in some conditions or diseases. Here, we present BANDITS, a R/Bioconductor package to perform differential splicing, at both gene and transcript-level, based on RNA-seq data. BANDITS uses a Bayesian hierarchical structure to explicitly model the variability between samples, and treats the transcript allocation of reads as latent variables. We perform an extensive benchmark across both simulated and experimental RNA-seq datasets, where BANDITS has extremely favorable performance with respect to the competitors considered.

## Background

Alternative splicing plays a fundamental role in the biodiversity of proteins as it allows a single gene to generate several transcripts and, hence, to code for multiple proteins [1]. However, variations in splicing patterns can be involved in development and disregulated in disease [2–4]. Differential splicing (DS) studies how splicing patterns vary between experimental conditions, and specifically, differential transcript usage (DTU) represents a primary branch to investigate DS [5]. DTU is present when there are changes, between two or more conditions, in the relative abundances of transcripts (i.e., in the transcript proportions), irrespective of the overall output of transcription. Alternative approaches to investigate DS are differential exon usage (DEU) [6], event specific differential splicing based on percent-spliced-in [7–9], and differential transcript expression (DTE) [5], which focuses on changes in the overall abundance of isoforms and, hence, identifies both differential gene expression (DGE) as well as differential splicing. Note that, although we broadly refer to differential splicing, all these approaches target differences in annotated transcripts (or exons), which may arise due to differential splicing as well as alternative start and terminal sites of the same transcript.

A significant challenge of DTU, and in general of DS, is that transcript-level counts (i.e., the number of RNA-seq reads originating from each isoform), which are of primary interest, are not observed because most reads map to multiple transcripts (and sometimes, multiple genes). Quantification tools [10] such as Salmon [11] or kallisto [12] allow, via expectation maximization (EM) algorithms, to estimate the expected number of fragments originating from each transcript. Most methods for DS (notably, DRIMSeq [13], BayesDRIMSeq [14] and SUPPA2^[1]^ [9]) follow a *plug-in* approach by inputing transcript estimated counts (TECs) and treating them as observed counts, thus neglecting the uncertainty in the estimates. In an attempt to mitigate this issue, rats [15] inputs TECs together with their bootstrap replicates; nevertheless rats is limited by the fact that it uses a G-test based on the Multinomial distribution, which assumes all biological replicates to share the same relative transcript abundance.

Instead of considering TECs, some methods, such as DEXSeq [6] and limma (via *diffSplice* function) [16], perform DEU by testing exon bin counts, which are observed directly; however, reads overlapping multiple exon bins are counted multiple times, once for each exon bin they map to. Furthermore, differential testing is done at the exon level, while transcript-level tests and proportions cannot be computed; for this reason, DEU is widely considered as a surrogate for DTU [5]. An alternative approach, ignoring the quantification step, considers the groups of transcripts that reads are compatible with, usually referred to as equivalence classes (ECs), and the respective counts. Recently, two articles [17, 18] proposed to perform DTU by applying DEXSeq on transcript estimated counts or on equivalence classes counts (ECCs); however, both approaches have limitations. The former, similarly to DRIMSeq, BayesDRIMSeq and SUPPA2, inputs TECs while ignoring their inherent variability. The latter, instead, has limited interpretability because testing cannot be done at the transcript level and transcript-level proportions cannot be computed; moreover, equivalence classes containing transcripts from distinct genes are excluded from the analyses. A further method considering ECs is cjBitSeq [14, 19], which performs a full Bayesian analysis and samples the allocation of each read to its transcripts of origin; however cjBitSeq, similarly to rats, does not allow for sample-specific proportions. Moreover, in the DTU implementation of cjBitSeq^[2]^, the equivalence classes containing transcripts from multiple genes are considered multiple times (once for each gene contained in the EC).

In order to overcome the limitations of current methods for DTU, we present BANDITS (Bayesian ANalysis of DIfferenTial Splicing), a R/Bioconductor package to perform DTU between two or more groups of samples, based on RNA-seq data. BANDITS uses a Bayesian hierarchical model, with a Dirichlet-multinomial structure, to explicitly model the sample-to-sample variability between biological replicates, which allows each sample to have distinct transcript relative abundances due to biological variability. BANDITS inputs the equivalence classes and respective read counts treats the transcript allocations of reads as latent variables, i.e., as parameters that are sampled, jointly with the model parameters, via Markov chain Monte Carlo (MCMC) techniques. In this way our method models the uncertainty arising from reads mapping to multiple transcripts (regardless of the origin of the annotated transcript, e.g., protein-coding gene, pseudogene, homologous gene, etc.). ECCs can be obtained by aligning reads either to a reference transcriptome, with pseudo-aligners Salmon [11] and kallisto [12], or to a reference genome with splice-aware genome aligner STAR [20], and computing the ECCs of the aligned reads via Salmon.

Despite the abundance of DS methods available in the literature, BANDITS introduces some unique features and, in both simulation and experimental data analyses, shows very favourable performance with respect to all the competitors we considered. Supplementary Table S1 summarizes the main features of the most popular methods for DTU based on RNA-seq data. BANDITS is the only DS tool that jointly allows for sample-specific proportions between biological replicates while also sampling the transcript allocation of reads. It is also the only DS method to sample the gene allocation of reads in equivalence classes that contain transcripts from distinct genes (Cmero et al. [18] exclude these ECs, while cjBitSeq considers these classes multiple times, once per gene). Furthermore, BANDITS is the first work to correct for the transcript (effective) lengths when computing the relative abundance of isoforms; hence, it is able to disentangle the probability that reads map to a transcript, from the probability of expressing a transcript (see Results), and uses the latter parameter for statistical testing. BANDITS tests for DTU at both transcript and gene level, allowing scientists to investigate what specific transcripts are differentially used (DU) in selected genes. Furthermore, our tool is not limited to two group comparisons and also allows to test for DTU when samples belong to more than two groups. Finally, despite the computational complexity of full MCMC algorithms, the MCMC sampling is coded in C++, which makes BANDITS highly efficient and feasible to run on a laptop, even for complex model organisms.

## Results

### The BANDITS hierarchical model

Consider a gene with *K* transcripts and *N* samples (i.e., biological replicates) from a given group. We define the latent vector of transcript-level counts for the *i*-th subject as 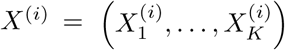, where 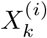 indicates the number of reads originating from the *k*-th transcript in the *i*-th sample, with *i* = 1, …, *N* and *k* = 1, …, *K*. We use a Bayesian hierarchical model [21, 22], which represents a natural approach to gather information from distinct samples, while allowing for sample-specific parameters, in a statistically rigorous way. We assume that *X*^(*i*)^ was generated from a multinomial distribution:

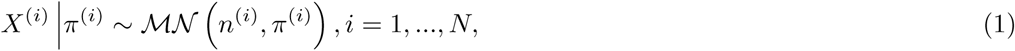

where 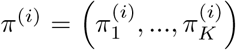, with 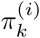 indicating the relative abundance of the *k*-th transcript within the gene in the *i*-th sample, *n*^(*i*)^ represents the total number of counts arising from the gene of interest in the *i*-th sample, and ℳ𝒩 (·) denotes the multinomial distribution. Assuming independence between genes, the full likelihood for all *N* samples in a group is defines as:

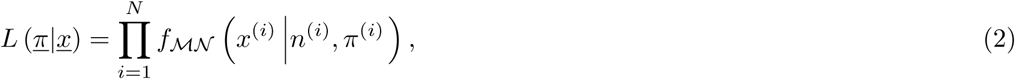

where *f*_*ℳ𝒩*_(·) indicates the density of the Multinomial distribution, *<u>π</u>* = (*π*^(1)^, …, *π*^(*N*)^), and *<u>x</u>* = (*x*^(1)^, …, *x*^(*N*)^), with 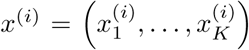 being the realization of the random variable *X*^(*i*)^, *i* = 1, …, *N*.

The transcript proportions for each sample are connected via a common Dirichlet prior distribution:

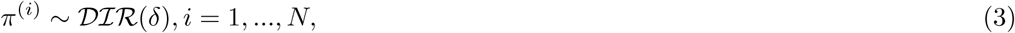

with 𝒟ℐℛ (·) denoting the Dirichlet distribution and *δ* = (*δ*_1_, …, *δ*_*K*_), where 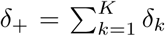 is the precision parameter, modelling the degree of over-dispersion between samples, and 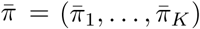, with 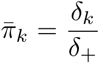 indicating the mean relative abundance of the *k*-th transcript, for *k* = 1, …, *K*. The prior distribution for the hierarchical parameters is:

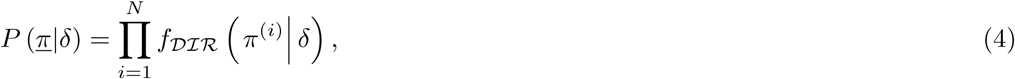

where *f*_𝒟ℐℛ_ (·) indicates the density of the Dirichlet distribution.

In order to exploit the information from other genes, we take advantage of DRIMSeq [13] to infer genewise precision parameters, and use these estimates to formulate an informative prior for *δ*_+_. If precision estimates are not computed, all *δ*_*k*_ parameters follow a vaguely informative prior distribution (see Methods).

Since most reads map to multiple transcripts, transcript-level counts are typically not observed directly. BANDITS inputs, for every gene, the equivalence classes of transcripts and respective counts, while the transcript-level counts are treated as latent variables and are sampled together with the model parameters (see Methods). In ECs with transcripts from more than 1 gene, the gene allocation of reads is also treated as a latent variable and sampled within the MCMC scheme (see Supplementary Section S1.2).

### MCMC overview

In order to infer the posterior distribution of the model parameters, we developed a Metropolis-within-Gibbs [23–25] MCMC algorithm where parameters are alternately sampled in three blocks: *δ*, via a Metropolis algorithm [24, 25] with an adaptive random walk proposal [26], *<u>π</u>* and *<u>X</u>*, both via a Gibbs sampler [27, 28]. The mathematical details of the sampling scheme are illustrated in Supplementary Section S1.1.

After discarding an initial *burn-in*, the convergence of chains and a potentially wider *burn-in* are assessed via Heidelberger and Welch’s stationarity test [29]. To avoid potential false positive results due to poor mixing, if the gene-level test has a p-value below 0.1, a second independent MCMC chain is run and results are recomputed on the aggregation of the two chains (*burn-in* excluded).

### Accounting for transcript lengths

We introduce a conceptual distinction between the probability that reads map to a transcript, which depends on the transcript length, and the probability that a gene expresses a transcript. While the former parameter is typically used to test for DTU, we argue that the latter should be employed instead, because it reflects the number of transcripts expressed by a gene, independently of their length. We use the mean relative abundance of transcripts, 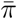, to compute the average probability of expressing transcripts, 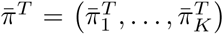, where 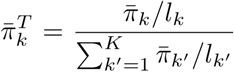, with *l*_*k*_ being the effective length of the *k*-th transcript, for *k* = 1, …, *K*. In the previous formula, at the numerator we normalize 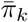 with respect to the effective length of the *k*-th isoform, while the denominator term is a scaling factor to ensure that 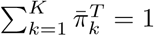. In three simulation studies, we compared results from BANDITS, which tests for DTU via 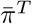, to a modified version of BANDITS using 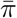. In all scenarios, and for both gene- and transcript-level tests, using 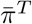 leads to improved performance (Supplementary Figures S25-S26). Furthermore, unlike other methods for DTU, BANDITS provides users an estimate of the actual mean transcript relative expression, 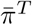.

### DTU testing

After inferring the model parameters, we test for DTU by comparing 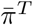 between conditions. Given groups *A* and *B*, with average transcript relative expression, for the *k*-th transcript, 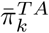 and 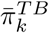, respectively, we test the following system of hypotheses:

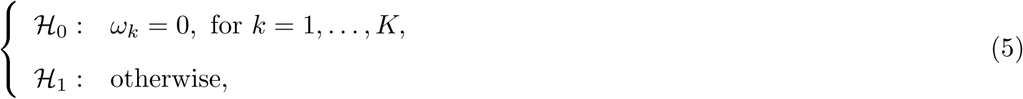

where 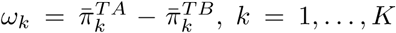. We approximate the posterior distribution of *ω* = (*ω*_1_, …, *ω*_*K*_) with a multivariate normal density [30], 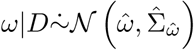, where 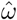 represents the posterior mode of *ω* and 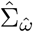 its covariance matrix, both inferred from the posterior chains, *D* denotes the input data (i.e., the ECCs) and 𝒩 (*µ*, Σ) indicates the normal density with mean *µ* and covariance Σ. In order to test for DTU at the gene level, BANDITS performs a multivariate Wald test [31], based on the normal approximation of *ω*, to test the set of hypotheses (5).

Our method also can unravel the specific transcripts that are DU by testing, for the *k*-th transcript, the following system of hypotheses: ℋ_0_ : *ω*_*k*_ = 0, vs. ℋ_1_ : *ω*_*k*_ ≠ 0. Similarly to the gene-level test, we perform a univariate Wald test based on the normal approximation of the marginal posterior distribution of 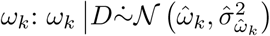, where 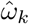 and 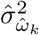 represent the posterior mode and variance of *ω*_*k*_, respectively, both inferred from the posterior chains. In both gene and transcript-level testing, false discovery rate (FDR) control is obtained by adjusting p-values via Benjamini-Hochberg correction [32].

BANDITS also outputs conservative gene and transcript-level scores, as well as a measure of the strength of DTU (see Methods). Furthermore, our method also allows to test for DTU between 3 or more conditions (see Supplementary Section S1.3).

### Simulation studies

We performed three RNA-seq stimulation studies to benchmark BANDITS against nine other DS methods.

First, we considered the human simulation from Soneson et al. [33], where two groups of 3 samples each are compared, and DU genes are simulated by inverting the relative abundance of the two most expressed transcripts across conditions.

By editing Soneson et al. [33] pipeline, we also built a second simulation dataset, from a human genome with two groups of 6 samples each, where DU genes are simulated by randomly permuting the relative abundance of the four most expressed transcripts; if a DU gene has two or three transcripts only, then their expression is permuted. In our view, this second simulation provides a more varied scenario compared to the first one: the dominant transcript (i.e., the most abundant isoform) does not always change between conditions and some genes will exhibit more changes, but whose magnitude might be smaller. This simulation is made available via FigShare (DOI *10*.*6084/m9*.*figshare*.*9467144, 10*.*6084/m9*.*figshare*.*9692429* and *10*.*6084/m9*.*figshare*.*9692918*). We will refer to the former and latter datasets as “3 vs. 3” and “6 vs. 6”, respectively. Further details about the simulations are reported in Methods, while a description of differential analyses and software versions can be found in Supplementary Section S1.4 and Table S2.

As a third scenario, we considered the 6 vs. 6 simulation and filtered transcripts, before the differential analyses, based on Salmon estimated counts: we kept transcripts with least 10 counts (across all samples) and an average relative abundance of at least 0.01.

We benchmarked BANDITS against several competitors: BayesDRIMSeq, cjBitSeq, DEXSeq, DEXSeq on ECCs (DEXSeq_ECCs), DEXSeq on TECs (DEXSeq_TECs), DRIMSeq, limma (via *diffSplice* function), rats and SUPPA2. We also consider the conservative gene and transcript-level scores from BANDITS, BANDITS_inv and BANDITS_maxGene (see Methods), as well as the ones from BayesDRIMSeq and cjBitSeq, that we call BayesDRIMSeq inv and cjBitSeq inv. Note that SUPPA2 does not perform a global gene-level test: in order to obtain a gene-level score we considered the minimum of the transcript-level adjusted p-values. For cjBitSeq transcript-level test, we used the probability that a transcript is not differentially used; note that this does not guarantee FDR control. Genes and transcripts with less than 20 and 10 estimated counts (across all samples), respectively, are excluded from Figures and Tables.

Figures 1 and 2 report the true positive rate (TPR) vs. FDR curves of all methods for gene and transcript-level tests, respectively. Note that fewer methods are displayed in transcript-level plots, because not all tools perform a transcript-level test. To facilitate graphical interpretation, for each method, we only report three dots corresponding to the observed FDR at 0.01, 0.05 and 0.1 thresholds; the full curves are available in Supplementary Figures S1 and S2. BANDITS exhibits highly favourable performance in all scenarios. In both, unfiltered and filtered, 6 vs. 6 simulation studies, BANDITS and its conservative scores (BANDITS_inv or BANDITS_maxGene) have the highest curves, while they are only second to SUPPA2 in the 3 vs. 3 simulated data. Furthermore, in all cases, BANDITS provides good control of the FDR, particularly for the 0.05 and 0.1 thresholds, while most methods show a significant deviation from these cut-offs. Compared to the original BANDITS tests, the conservative scores, BAN-DITS_inv and BANDITS_maxGene, provide a better FDR control without lowering the overall curve. Note that in the 3 vs. 3 simulation, BANDITS_inv, BayesDRIMSeq_inv and cjBitSeq_inv scores are favoured by the fact that DTU genes are simulated by inverting the two most expressed transcripts, hence the dominant transcript always changes between conditions in DU genes.

**Figure 1.**
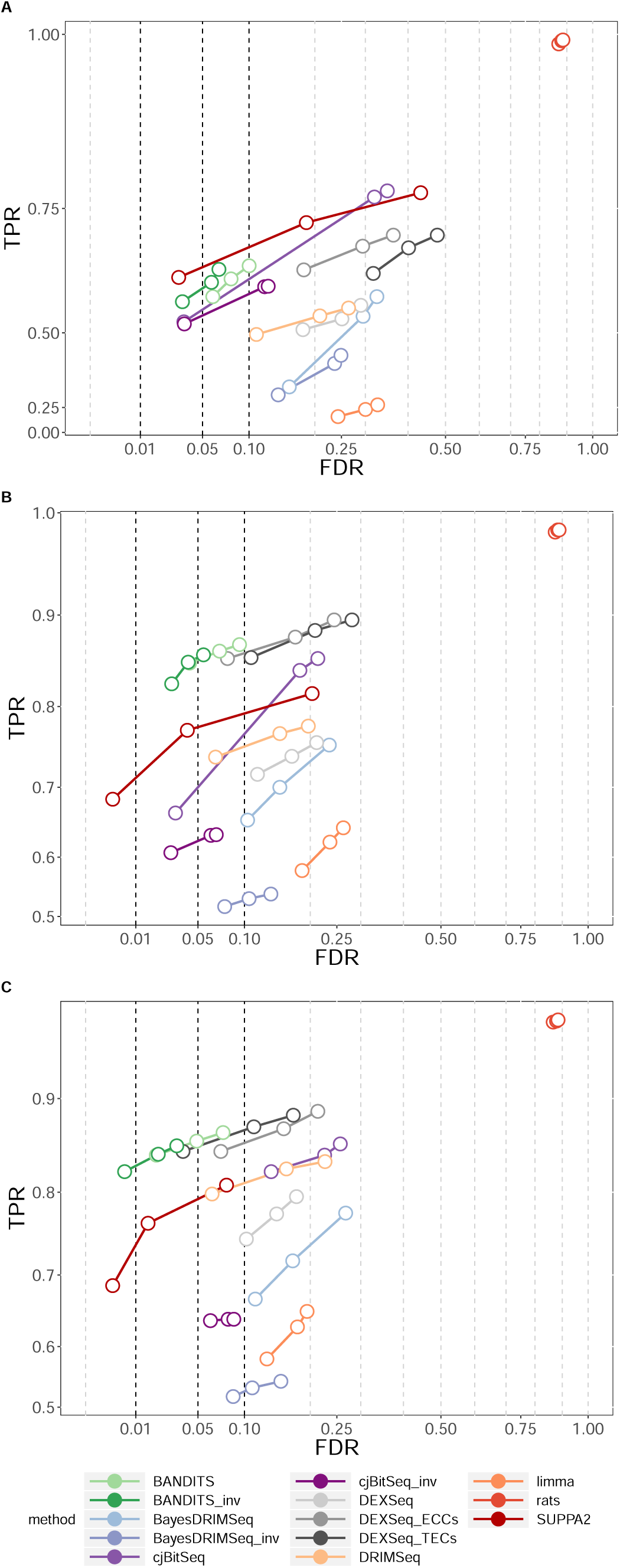
TPR vs. FDR for gene-level testing for the three 2-group comparison simulation studies. A) 3 vs. 3 simulation study; B) 6 vs. 6 simulation study; C) 6 vs. 6 simulation study with transcript pre-filtering (transcripts with at least 10 counts and an average relative abundance of 0.01). Circles indicate observed FDR for 0.01, 0.05 and 0.1 significance thresholds.

**Figure 2.**
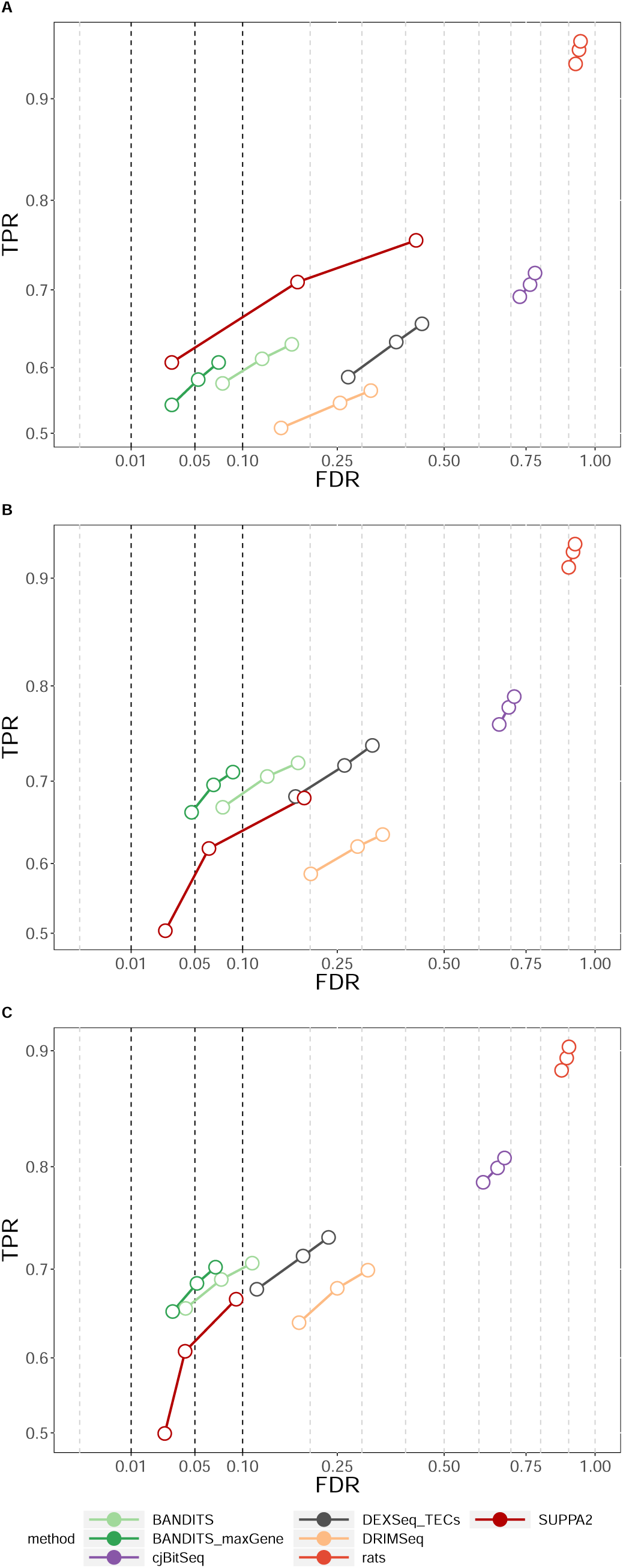
TPR vs. FDR for transcript-level testing for the three 2-group comparison simulation studies. A) 3 vs. 3 simulation study; B) 6 vs. 6 simulation study; C) 6 vs. 6 simulation study with transcript pre-filtering (transcripts with at least 10 counts and an average relative abundance of 0.01). Circles indicate observed FDR for 0.01, 0.05 and 0.1 significance thresholds. Note that, for cjBitSeq, we considered the probability that a transcript is not differentially used, which does not guarantee FDR control.

Supplementary Figures S3 and S4 compare results obtained by BANDITS, in both 3 vs. 3 and 6 vs. 6 simulated data, on the original data and when filtering lowly abundant transcripts: in both cases, and particularly in the 3 vs. 3 simulation, transcript pre-filtering leads to an improvement of gene and transcript-level testing.

To show the ability of BANDITS to compare more than 2 groups, we performed a DTU analysis between 3 groups. We considered the 6 vs. 6 simulation and separated the first group in two sub-groups of size 3, hence obtaining a structure with 3 groups of size 3, 3 and 6. Again, we repeated the differential analyses with and without transcript pre-filtering (minimum 10 counts and an average relative abundance of 0.01 per transcript). TPR vs. FDR curves are visible in Supplementary Figures S10-S11: BANDITS again has favourable performance in both scenarios, particularly when pre-filtering transcripts. Note that only BANDITS, DRIMSeq and DEXSeq variants are plotted because they are the only methods considered in this work that allow to jointly compare more than two groups (see Supplementary Table S1).

### Experimental data analyses

We also applied the previous DTU models to two RNA-seq experimental datasets. First, we studied the human data from Best et al. [9, 34], consisting of a two group comparison with 3 samples in each group, where 83 splicing events, corresponding to 82 genes, were validated via reverse transcriptase polymerase chain reaction (RT-PCR). We restricted our study to the most 10,000 expressed genes (given Salmon estimated counts), which include all 82 validated genes. We will refer to this database as “Best et al.”.

Figure 3 shows the receiving operating characteristic (ROC) curves of all methods considered for gene-level testing, while Table 1 reports the area under the curve (AUC), the partial AUC of levels 0.1 and 0.2, the median position of the 82 validated genes in the raking of 10,000 analyzed genes, and the number of validated genes amongst the most significant 100 and 200 genes for each method. BANDITS has again very favourable performance: BANDITS and BANDITS_inv provide the two lowest median rankings for the validated genes, as well as the highest (overall and partial) AUCs, and the highest TPR curves for false positive rate (FPR) between 0 and 0.25; furthermore, BANDITS top ranked genes contain more validated genes than any other method.

**Table 1.** Results from the “Best et al.” experimental dataset; methods are sorted by lowest “Median position”. “Median position” indicates the median position of the 83 validated genes in the ranking of 10,000 analyzed genes; AUC refers to the area under the ROC curve; pAUC 0.1 and 0.2 represent the partial AUC of levels 0.1 and 0.2, respectively; “top 100” and “top 200” report the number of validated genes (82 in total) in the 100 and 200 genes with lowest FDR from each method; “GO 0.01” and “GO 0.05” indicate the fraction of “validated GO terms” found by each method, when considering FDR thresholds 0.01 and 0.05, respectively.

|  | Median<br>position | AUC | pAUC<br>0.1 | pAUC<br>0.2 | top<br>100 | top<br>200 | GO<br>0.01 | GO<br>0.05 |
| --- | --- | --- | --- | --- | --- | --- | --- | --- |
| BANDITS_inv | 596 | 0.81 | 0.04 | 0.11 | 16 | 24 | 0.32 | 0.35 |
| BANDITS | 673 | 0.80 | 0.04 | 0.11 | 18 | 25 | 0.34 | 0.33 |
| cjBitSeq | 900 | 0.79 | 0.04 | 0.10 | 9 | 20 | 0.15 | 0.34 |
| rats | 942 | 0.80 | 0.03 | 0.10 | 10 | 16 | 0.34 | 0.34 |
| DEXSeq_TECs | 968 | 0.79 | 0.03 | 0.09 | 11 | 20 | 0.33 | 0.30 |
| DEXSeq_ECCs | 1039 | 0.78 | 0.03 | 0.10 | 12 | 20 | 0.30 | 0.29 |
| BayesDRIMSeq | 1231 | 0.74 | 0.02 | 0.08 | 6 | 9 | 0.24 | 0.27 |
| DEXSeq | 1348 | 0.78 | 0.03 | 0.08 | 7 | 12 | 0.26 | 0.29 |
| limma | 1556 | 0.74 | 0.03 | 0.08 | 9 | 13 | 0.33 | 0.29 |
| SUPPA2 | 2110 | 0.67 | 0.02 | 0.07 | 11 | 13 | 0.22 | 0.27 |
| DRIMSeq | 3248 | 0.59 | 0.03 | 0.07 | 14 | 20 | 0.30 | 0.27 |
| cjBitSeq_inv | 5146 | 0.59 | 0.02 | 0.05 | 12 | 15 | 0.20 | 0.35 |
| BayesDRIMSeq_inv | 5362 | 0.57 | 0.02 | 0.04 | 3 | 10 | 0.14 | 0.18 |

**Figure 3.**
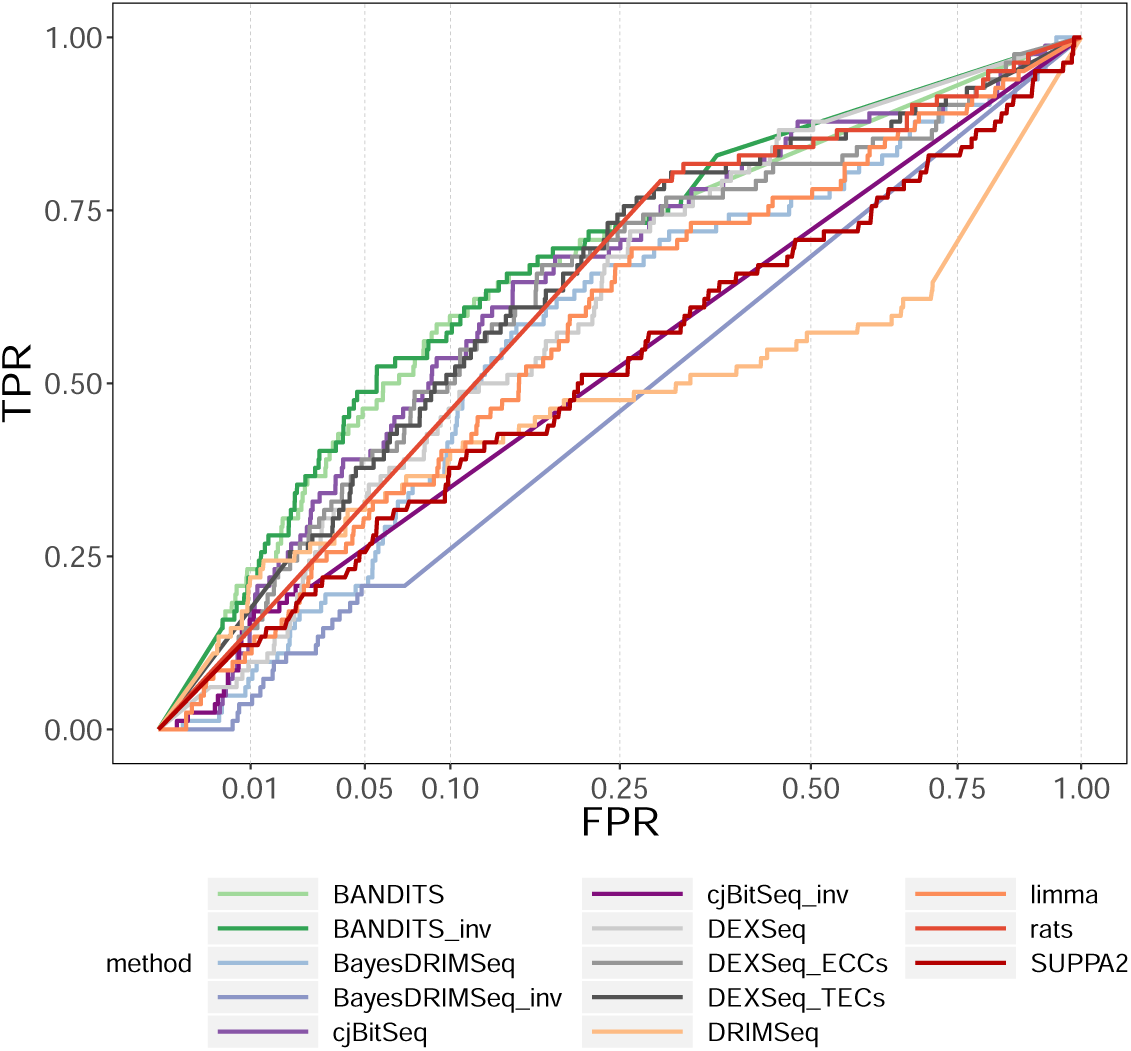
ROC curve (TPR vs. FPR) for gene-level testing in the “Best et al.” experimental dataset.

We also performed a gene ontology (GO) analysis based on the 82 validated genes, and obtained 338 (out 22,911) enriched GO terms with a p-value below 0.01, which we define as our “validated GO set”. We then inferred enriched GO terms based on the significant genes from each method, i.e., with FDR below 0.01 and 0.05, and selected the most significant 338 GO terms. The fractions of terms in the “validated GO set” captured by BANDITS and BAN-DITS_inv are among the highest and vary between 0.32 and 0.35, while, for the other methods, this value ranges between 0.14 and 0.35 (Table 1).

Moreover, to add biological perspective, in Supplementary Section S1.5, Tables S9-S10 and Figures S13-S17 we present an in depth visual inspection of two genes, and show how BANDITS can be used to accurately gain novel biological insight by identifying differentially spliced genes, as well as the individual transcripts that are affected.

We further considered a second human experimental dataset [35]. Here, we performed a “null” analysis to investigate FPRs, by comparing two groups of 3 healthy patients each. Again, we removed genes with less than 20 estimated counts across all samples.

Figure 4 shows the gene-level test FPR vs. FDR curves of each method. Supplementary Figures S7 and S8 report the same analysis for both gene and transcript-level tests, when considering raw and adjusted p-values, while Supplementary Table S4 displays the FPRs obtained at the 0.05 threshold. Overall limma, BANDITS, BANDITS_inv, DRIMSeq and DEXSeq display the lowest FPRs at the gene level; BANDITS BANDITS_maxGene and DRIMSeq also lead to the lowest FPRs when considering transcript-level tests. Instead, rats, DEXSeq_ECCs and DEXSeq_TECs provide the worst control of FPs in gene-level tests, particularly for 0.01 and 0.05 thresholds, while rats has the highest number of false positives when testing transcript.

**Figure 4.**
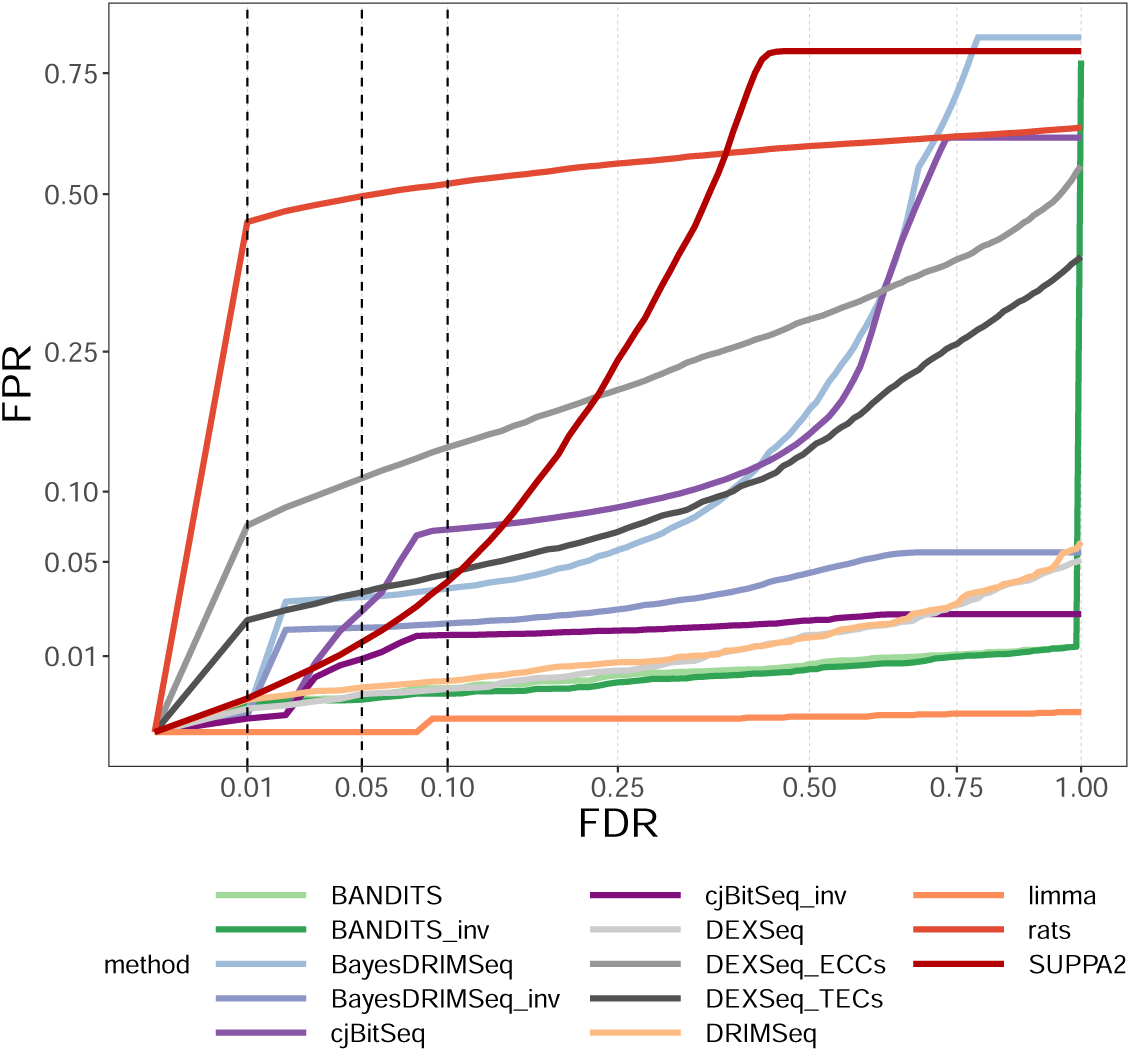
FPR vs. FDR for gene-level testing in the null experimental dataset.

### Computational benchmark

We performed a computational comparison of all the methods considered in the 6 vs. 6 simulation study, with and without transcript pre-filtering. Analyses were run on 12 cores, when parallelization was allowed, on our Opteron 6100 server.

Figure 5 and Supplementary Tables S6 and S7 illustrate the computational cost of each method. In our benchmark, cjBitSeq stands out as the most computationally intensive tool, both in the alignment (via Bowtie2) and differential components, followed by DEXSeq and limma, mostly due to the python *dexseq_count*.*py* function which translates the genomic alignments of reads into exon bin counts. On the opposite side DEXSeq_TECs and DRIMSeq, which use transcript estimated counts, are the fastest methods to run. Overall, BANDITS is significantly faster than cjBitSeq, DEXSeq and limma, but slower than DEXSeq_ECCs and than tools using TECs; nonetheless, BANDITS has a 3 time speed-up when pre-filtering transcripts, bringing it close to DEXSeq_ECCs. Considering this significant computational gain, and the improved performance obtained when pre-filtering transcripts (Supplementary Figures S3 and S4), we highly encourage users to filter lowly abundant transcripts, which can be done automatically in BANDITS via the *filter_transcripts* function. Furthermore, we found that BANDITS scales well when increasing sample size: it required 43.5 minutes when using 2 samples per group, 50.5 with 3 and 58.8 with 6 (details in Methods).

**Figure 5.**
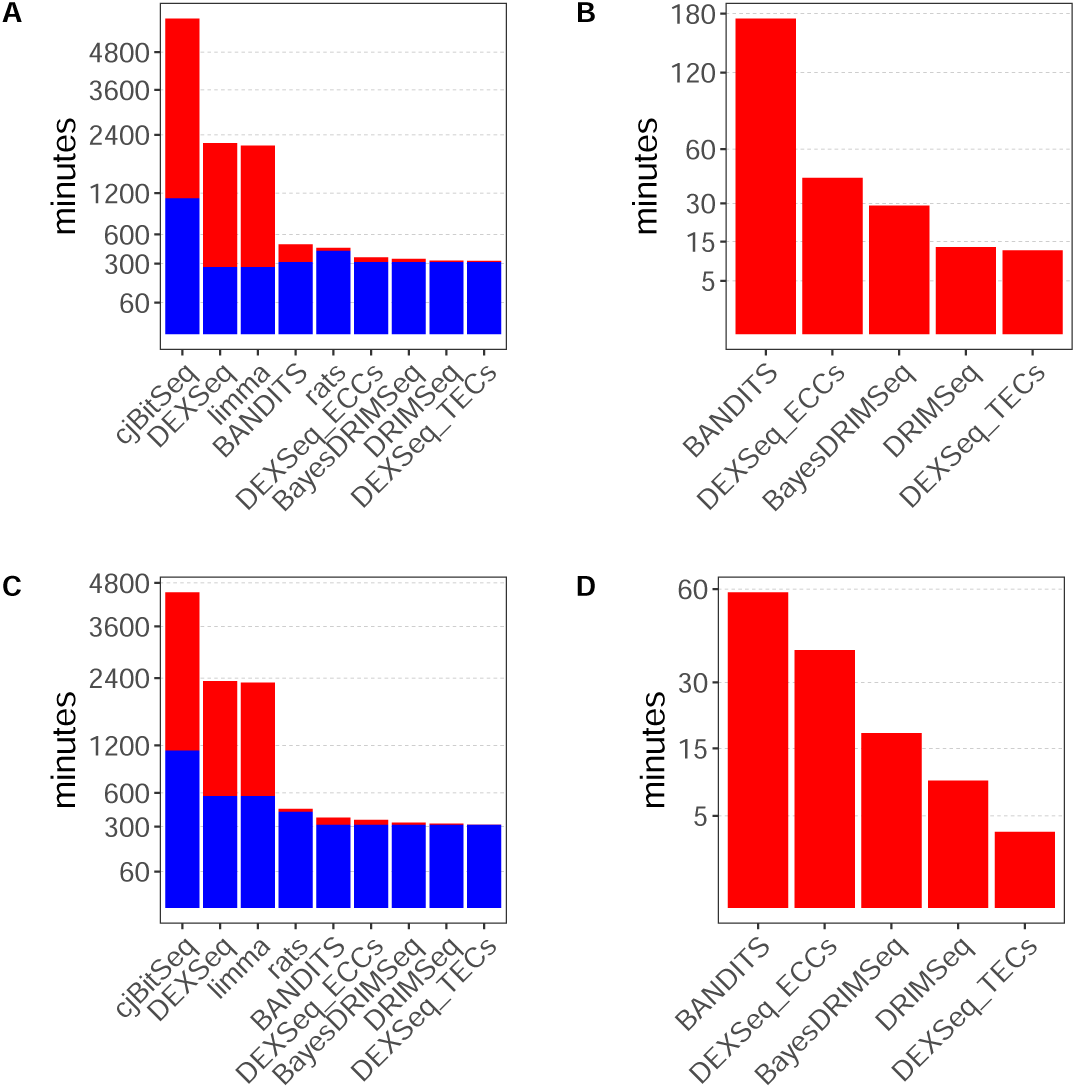
Computational benchmark in the 6 vs 6 simulation study. the (computational) cost of alignment and quantification (when required) is shown in blue; the cost of the differential analyses is shown in red. A) Full pipeline; B) differential analyses after running STAR and Salmon; C) full pipeline, when filtering the transcriptome; D) differential analyses after running STAR and Salmon, when filtering the transcriptome. cjBitSeq, DEXSeq, limma and rats are excluded from B) and D) because they require a distinct alignment pipeline. Details in Supplementary Section S1.4.

Note that, except cjBitSeq, DEXSeq and limma, the cost of alignment (via STAR) and quantification (via Salmon) is much higher than the cost of the differential analyses, making the overall cost of the full pipelines of these methods similar.

Supplementary Table S8 and Figure S12 also report the maximum RAM used by each method. The alignment step (via STAR) required significantly more memory than any differential method, with a maximum usage of more than 30 gigabytes (GBs). On both unfiltered and filtered transcriptomes, DEXSeq was the differential method with the highest maximum RAM usage, with 10.2 and 5.2 GBs of RAM needed, while BANDITS showed modest memory usage and required at most 1.8 and 0.9 GBs.

### Stratification by expression level

To investigate how method performance is influenced by gene abundance, we stratified the results of the 6 vs. 6 simulation study, and of both experimental data analyses according to gene expression, by grouping genes into lowly (first tertile), medium (second tertile) and highly expressed (third tertile).

In the simulation study (Supplementary Figure S5), the ordering of methods is roughly unaltered, while medium and highly expressed genes have a general better FDR control compared to lowly abundant ones. In the Best et al. data analysis (Supplementary Figure S6 and Table S3), medium and highly expressed genes tend to have a better ranking (e.g., median position of validated genes) compared to lowly abundant ones, but no method outperforms the others in all three cases. Finally, the null data analysis (Supplementary Figure S9 and Table S5) shows that more genes are erroneously detected as their expression increases; in particular, rats and DEXSeq_ECCs show worrying FPRs of 82.65% and 29.73%, respectively, for highly expressed genes, given an FDR significance threshold of 0.05. BANDITS and BANDITS_inv, instead, provide among the lowest false detections in any group of genes, with FPRs ranging between 0.05% and 0.42%.

### Sensitivity analyses

We performed two sensitivity analyses to investigate how robust BANDITS results are to the prior specification and to the alignment and quantification tools.

Firstly, we investigated how sensitive BANDITS is to the prior used for the precision parameter. We ran all simulation and experimental data analyses with vaguely informative prior for the precision parameter, i.e., with BANDITS default prior when no informative prior is provided. BANDITS performance marginally decreases when no informative prior is supplied, particularly concerning FDR control, even though pre-filtering transcripts alleviates this issue and leads to more similar results (Supplementary Tables S11-S12 and Figures S18-S22).

Secondly, we studied how results were affected by alignment and quantification tools. We considered the 6 vs. 6 simulation study and aligned and quantified reads with transcript pseudo-aligners Salmon [11] and kallisto [12], and with splice-aware genome aligner STAR [20] (transcript abundances estimated with Salmon on the aligned reads). We ran BANDITS, with and without transcript pre-filtering, on the three input data. BANDITS results originating from Salmon and STAR were fairly similar, while kallisto lead to higher FDR for approximately the same TPR, particularly for gene-level testing (Supplementary Figures S23-S24).

## Discussion

In this manuscript, we have introduced a method to perform differential splicing based on RNA-seq data. BANDITS uses a Bayesian hierarchical structure to model the variability between samples, and treats the transcript (and gene) allocations of reads as latent variables; model parameters and latent variables are sampled via MCMC techniques. We designed benchmarks, based on three simulation studies and two experimental data analyses, where we compared BANDITS against the most popular methods for differential splicing. Results highlight BANDITS strong performance, and provide a comprehensive guide for users interested in choosing a tool to investigate DS.

A limitation in common to all methods considered, is to rely on an annotated transcriptome (and genome, for genome alignment), which may lead to inaccurate inference in case of misannotated transcripts and genes [36]; this phenomenon might be particularly present for disease samples, whose condition might lead to the development of unannotated transcripts or genes (e.g., gene fusions). Therefore, all DS methods considered here would benefit from the development of tools that enhance the annotated transcriptome based on the available data, hence accounting for the particular features of the samples considered. Furthermore, BANDITS targets genes and transcripts undergoing differential splicing, but does not identify specific splicing events. Some tools, most notably SUPPA2, target local splicing events (e.g., intron retention or exon skipping), usually based on percent-spliced-in. However, such an approach typically leads to lower power than jointly considering all reads available for a gene. A further limitation of BANDITS is that it does not allow for covariates; to overcome this issue, we introduced a regression structure in our model to incorporate covariates, such as batches. However, when adding batch effects to our simulation studies, even in extreme scenarios, the original version of BANDITS outperformed, in terms of power and FDR, the modified version allowing for covariates (data not shown). Moreover, we noticed that BANDITS was very robust to batch effects, which only marginally altered its performance. This suggests that the misspecification of the model (i.e., ignoring batches when present) might be less deleterious than having a more complex modelling structure, involving more parameters. Therefore, we choose not to include this modification in the final version of BANDITS.

Finally, we note that BANDITS, although developed with a focus on RNA-seq data, can also be applied to long-read sequencing data. Soneson et al. (2019) [36] found that Illumina RNA-seq reads and Oxford Nanopore Technologies long reads generated equivalence classes with almost equivalent average number of transcripts. Hence, one might expect at least the current generation of long read transcriptome data to also benefit from BANDITS transcript latent variable allocation approach.

## Conclusions

We presented BANDITS, a novel Bayesian method to investigate differential splicing from RNA-seq data. At present, our tool is the only method that jointly models the variability between biological replicates, by allowing for sample-specific proportions, and the mapping uncertainty of reads, by sampling their transcript (and gene) allocations. BANDITS is also the first DS tool to correct for the transcript effective lengths, allowing it to recover the actual probability of expressing a transcript. Our method tests, both, genes and transcripts for DS, and allows comparisons between more than two groups. We also introduce a measure of the DTU strength, which can be used as an alternative way to rank genes.

In all simulation and experimental datasets analyzed, BANDITS has extremely favourable performance and exhibits good FDR and excellent FPR control. Furthermore, despite requiring full MCMC inference, it is computationally competitive, particularly after applying reasonable expression level filters.

Finally, BANDITS is released as a R/Bioconductor package, which makes it easy to update, distribute and integrate within existing data analysis pipelines.

## Methods

### Prior distributions

Since the Dirichlet parameters *δ*_1_, …, *δ*_*K*_ are positive, we sample them and formulate their prior in the logarithmic scale, a common choice to improve mixing of positive parameters.

If gene-wise precision parameters are not computed (via *prior_precision* function), we specify a vaguely informative prior distribution for the logarithm of the Dirichlet parameters: *log*(*δ*_*k*_) ∼ 𝒩(*µ* = 0, *σ*^2^ = 100), *k* = 1, …, *K*.

Instead, if gene-wise precision parameters are available, we compute the mean and variance of their logarithm, 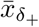 and 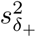, and formulate an informative prior for *log*(*δ*_+_) as: 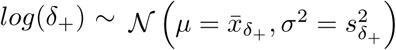. The remaining *K* − 1 Dirichlet parameters *a priori* are distributed as follows:

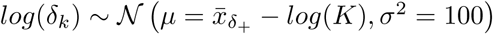, for *k* = 1, …, *K* −1, which corresponds to a vaguely informative prior; setting 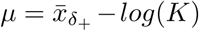 instead of 0, corresponds to assuming that, *a priori, δ*_+_ is equally distributed across the *K* transcripts. In order to obtain the prior distribution for *log*(*δ*_*K*_) we apply the change of variable via the Jacobian transformation ([37]).

### Latent variables allocation

We define the set of *J* equivalence classes available for a given gene as *C* = (*C*_1_, …, *C*_*J*_), where *C*_*j*_ indicates the list of transcripts present in the *j*-th equivalence class. Note that ECs not supported by any read are not included in *C*. The number of reads compatible with *C*_*j*_ in the *i*-th sample is denoted by 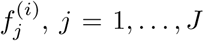 and *i* = 1, …, *N*. For ECs with at least two transcripts, reads in 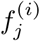 need to be allocated to the transcripts in *C*_*j*_. We introduce the vector 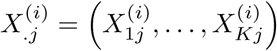, where 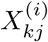 indicates the number of reads from the *j*-th EC that were generated from the *k*-th transcript in the *i*-th sample, with *j* = 1, …, *J, k* = 1, …, *K* and *i* = 1, …, *N*. Note that 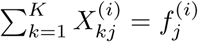 and 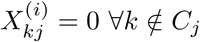.

Clearly, 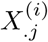 cannot be observed directly; it is hence treated as a latent variable which, under the assumption of uniform coverage, is sampled from the following density:

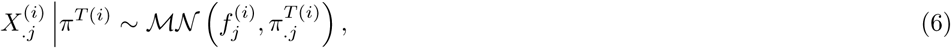

where 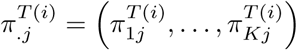, with

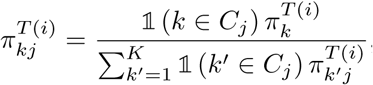, where 𝟙(a) is 1 if *a* is true, and 0 otherwise. Intuitively, 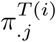 modifies *π*^*T*(*i*)^ to ensure that reads are only allocated to the transcripts in *C*_*j*_.

Once EC reads have been allocated to the respective transcripts, we can compute the corresponding counts for the *k*-th transcript by adding counts across 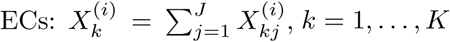 and *i* = 1, …, *N*.

If an equivalence class has transcripts from more than one gene, the probability vector 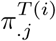 is modified to include all transcripts from the genes in the EC, and the transcript-level probabilities are weighted by the number of reads associated to each gene (details in Supplementary Section S1.2).

### Convergence diagnostic

BANDITS users can specify an initial number of iterations to discard as *burn-in* (minimum 2, 000), as well as the number of iterations the MCMC is run for after the initial *burn-in* (minimum 10, 000).

To ensure the posterior chains have reached convergence, after discarding the pre-specified *burn-in*, BANDITS performs Heidelberger and Welch (HW) stationarity test [29] on the marginal log-posterior of the hyper-parameters, i.e., *log*(*P*(*δ*|*π*)) ∝ *log*(*P*(*π*|*δ*)) + *log*(*P*(*δ*)); by adding the log-posterior densities from all groups, and performing a global convergence diagnostic test. A wider *burn-in* is removed, if estimated via HW test; moreover, if HW stationarity test is rejected at the 0.01 significance threshold, the full MCMC output is discarded and the algorithm is run again (up to three times).

Furthermore, when a gene-level test has a p-value below 0.1, BANDITS runs a second MCMC chain and, after removing the *burn-in*, recomputes the outputs based on the aggregation of the two chains.

### DTU test

For every gene, we test the system of hypothesis (5): since the *K* equations are linearly dependent, we only need to test *K* − 1 parameters; hence, we rewrite the system of hypothesis as:

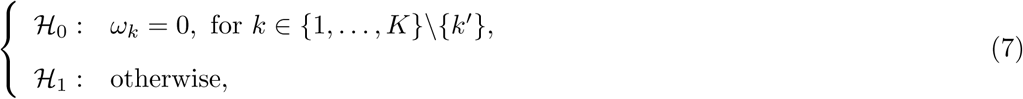

where *k*′ ∈ {1, …, *K*} is the transcript that should be removed from the test. The null distribution of *ω*_−*k*′_ = (*ω*_1_, …, *ω*_*k*′ −1_, *ω*_*k*′ +1_, … *ω*_*K*_) is approximately normal [30], with mean 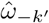 and covariance matrix 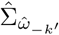, both inferred from the posterior chains. This leads to a multivariate Wald test [31] based on the null distribution of 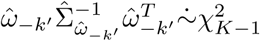, where 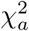 denotes the chi-square random variable with *a* degrees of freedom, and *b*^*T*^ and *b*^−1^ indicate the transpose and inverse of *b*, respectively. In order to choose the transcript to remove from the test, *k*′, we considered several options: randomly drawing one of the *K* transcripts, the transcript with the smallest expression, the isoform with the smallest difference between conditions, and averaging the p-values obtained from all *K* possible choices of *k*^1^. After benchmarking all four approaches, we choose the last one, because in our simulation studies it provided the highest sensitivity and best FDR control (data not shown).

Similarly, we test for differential usage in individual isoforms, by considering the system of hypothesis for the *k*-th transcript: ℋ_0_ : *ω*_*k*_ = 0 vs. ℋ_1_ : *ω*_*k*_ ≠ 0. In this case we use a univariate Wald test based on the statistic 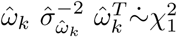, where 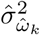 is the estimated marginal variance of *ω*_*k*_, inferred from the posterior chains.

Supplementary Section S1.3 shows how to extend this scenario when comparing 3 or more experimental conditions.

### Conservative scores and DTU measure

We propose two conservative scores for gene and transcript-level testing. The former is inspired by work from Papastamoulis and Rattray (2017) [14], where the authors propose to filter *a posteriori* all genes whose estimated dominant transcript (i.e., the most expressed transcript) is unchanged between conditions, leading to scores BayesDRIMSeq_inv and cjBitSeq_inv. How-ever, excluding all such genes, regardless of their significance, is an excessive filter in our opinion because genes might exhibit DS while preserving their dominant transcript. Here, when testing genes in two group comparisons, we introduce a moderated version of that score, that we call BANDITS_inv: we propose to inflate the adjusted p-value, defined as 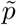, by taking its square root when the dominant transcript is unchanged between conditions. If the dominant transcript is estimated to change between conditions (according to the posterior mode of 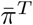), then 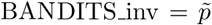, otherwise 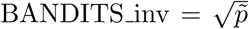. We further propose a conservative transcript score, called BANDITS_maxGene, which takes the maximum between the transcript and gene-level adjusted p-values; in this way, a transcript can only be selected if the corresponding gene is also significant.

Note that, in the Best et al. experimental data analysis, 41% of the validated genes are inferred to have distinct dominant isoforms between conditions, while this value decreases to 17% when considering non-validated genes; this fact seems to empirically justify our intuition of moderating Papastamoulis and Rattray’s inversion criterion.

For two group comparisons, we also propose a score, called *DTU_measure*, to measure the intensity of the differential usage change between conditions, similarly to fold changes in differential expression analyses. Given a gene with K transcripts and estimated mean relative transcript abundance 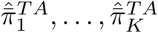, for group *A*, and 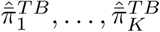, for group *B, DTU_measure* is defined as the summation of the absolute difference between the two most expressed transcripts: 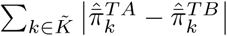, where 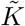 indicates the set of two most expressed transcripts across both groups. This measure ranges between 0, when proportions are identical between groups, and 2, when an isoform is always expressed in group *A* and a different transcript is always chosen in group *B*.

### Simulation

The 3 vs. 3 simulated data is taken from Soneson et al. [33]; when simulating reads for the 6 vs. 6 simulation study we modified the pipeline from Soneson et al. [33] to allow 6 samples per group.

In both cases, reads were simulated via RSEM [10]. using the Ensembl GRCh37.71 catalogue. First, RSEM (via *rsem-calculate-expression* module) was used on the human sample SRR493366 (http://www.ebi.ac.uk/ena/data/view/SRR493366) to estimate transcripts per million (TPM), which were then scaled to have 40 million reads for each sample and, via a mean-dispersion relationship derived from real data [38], used to obtain mean and dispersion parameters for each gene. Gene-level counts were simulated using these parameters via a negative binomial model, and then distributed to transcripts via a Dirichlet-multinomial distribution (with mean transcript-relative abundance parameters set to the isoform fractions estimated by RSEM on sample SRR493366).

Amongst genes with expected gene count above 500 and at least two transcripts with relative abundance above 10%, 1,000 genes were randomly selected to be differentially spliced. In the 3 vs. 3 simulation, differentially spliced genes were simulated by inverting, between groups, the relative abundance of the two most expressed transcripts. In the 6 vs. 6 simulation instead, differentially spliced genes were simulated by randomly permuting, in one group, the relative abundance of the four most expressed transcripts (if a gene had two or three transcripts only, then their expression was permuted).

Finally, the simulated transcript counts were used as input to RSEM to simulate, via *rsem-simulate-reads* module, 101 bp paired-ended fastq files for each sample.

### Scalability

We performed a computational benchmark of BANDITS, based on the 6 vs. 6 simulation, to investigate how computational times scale with respect to the sample size. We selected 2 and 3 samples per group and ran BANDITS on a 2 vs. 2 and 3 vs. 3 group comparison. In all cases, 12 cores from our Opteron 6100 based server were used, and the same transcripts were pre-filtered, based on the transcripts selected from the 6 vs. 6 analysis.

The computational cost scales less than linearly as the sample size increases: BANDITS took 43.5 minutes when using 2 samples per group, 50.5 with 3 and 58.8 with 6.

## Additional Files

Additional file 1: Methodological details, Additional Tables and Figures.

## Availability of data and materials

BANDITS is available on the Bioconductor site (https://bioconductor.org/packages/BANDITS) and on GitHub (https://github.com/SimoneTiberi/BANDITS).

For the 6 vs. 6 simulated data we designed, the fastq files for the 12 samples, TECs, ECCs and truth table are available at FigShare (https://figshare.com/projects/RNA-seq simulated data for differential transcript usage DTU analyses/66275), with DOI *10*.*6084/m9*.*figshare*.*9467144, 10*.*6084/m9*.*figshare*.*9692429* and *10*.*6084/m9*.*figshare*.*9692918*.

The code for simulating the RNA-seq reads and to perform all analyses is made available on GitHub (https://github.com/SimoneTiberi/BANDITSmanuscript) and on zenondo with DOI *10*.*5281/zenodo*.*3664468*.

The “Best et al.” [34] and “null” [35] experimental datasets are accessible at the Gene Expression Omnibus (GEO) under accessions GSE59335 and GSE37765, respectively.

## Author’s contributions

ST conceived the method, implemented it and performed the analyses. ST and MDR designed the study and wrote the manuscript. All authors read and approved the final article.

## Acknowledgements

The authors would like to thank Charlotte Soneson, Izaskun Mallona, Helena Lucia Crowell, Panagiotis Papastamoulis, Magnus Rattray, David Rossell and all members of the Robinson Lab for their helpful comments, suggestions and contributions to this work.

## Consent for publication

Not applicable.

## Ethics approval and consent to participate

Not applicable.

## Footnotes

[1] SUPPA2 performs both event-specific DS as well as canonical (transcript-level) DTU. Here, we only consider the DTU application of SUPPA2.

[2] cjBitSeq can perform both DTE and DTU analyses. Here, we refer to its DTU method only.

