## Additional file 1 for "BANDITS: Bayesian differential splicing accounting for sample-to-sample variability and mapping uncertainty"

**S2 Additional Tables and Figures** 7

#### S1 Methodological details

##### S1.1 MCMC sampling

This Section follows the notation introduced in the main manuscript.

Posterior chains are sampled via a Metropolis-within-Gibbs [1–3] Markov chain Monte Carlo (MCMC) algorithm. In each iteration, we alternately sample the model parameters from their conditional distributions, as shown below. Parameters from distinct experimental conditions (i.e., groups) are inferred separately.

The hyperparameters $\delta = (\delta_1, \ldots, \delta_K)$ are sampled, after applying the logarithmic transformation, from a Metropolis algorithm [2, 3] targeting the conditional distribution $\delta|\underline{\pi}$; proposal values are sampled from an adaptive random walk (ARW) scheme [4]. The sample-specific transcript proportions, $\underline{\pi} = \left(\pi^{(1)}, \ldots, \pi^{(N)}\right)$, are sampled, via a Gibbs sampler [5, 6], from their conditional distribution $\underline{\pi}|\delta, \underline{X}$. Similarly, the latent states, representing the unobserved transcript level counts, are sampled via Gibbs sampler from their conditional distribution $\underline{X}|\underline{\pi}, \underline{D}$, with $\underline{D} = \left(D^{(1)}, \ldots, D^{(N)}\right)$, where $D^{(i)}$ denotes the input data for the $i$-th sample (i.e., the set of equivalence classes counts).

We add a pre-subscript to all parameters, to indicate the value at the current iteration of the MCMC. We initialize the hyper and hierachical parameters as follows: ${}_0\delta_k = 1$ and ${}_0\pi_k^{(i)} = 1/K$, for $k = 1, \ldots, K$ and $i = 1, \ldots, N$. Note that the latent variables are not initialized because they are sampled from a Gibbs step, which does not require the value of the previous iteration. After initialising parameters, we update them according to the following scheme for $R$ iterations.

For $r = 1, \ldots, R$:

Update $\underline{X}|\underline{\pi}, \underline{D}$: For $i = 1, \ldots, N$, we performs steps I) and II) below.

I) First, for $j = 1, \ldots, J$, we sample the allocation of the $j$-th EC counts, $f_j^{(i)}$, to the $K$ transcripts as follows:

$$ {}_rX_{.j}^{(i)} | {}_{r-1}\pi^{T(i)}, f_j^{(i)} \sim \mathcal{MN}(f_j^{(i)}, {}_{r-1}\pi_{.j}^{T(i)}), \tag{S1} $$

where ${}_{r-1}\pi_{.j}^{T(i)} = \left( {}_{r-1}\pi_{1j}^{T(i)}, \ldots, {}_{r-1}\pi_{Kj}^{T(i)} \right)$, with ${}_{r-1}\pi_{kj}^{T(i)} = \dfrac{\mathbb{1}\left(k \in C_j\right) {}_{r-1}\pi_k^{T(i)}}{\sum_{k'=1}^K \mathbb{1}\left(k' \in C_j\right) {}_{r-1}\pi_{k'j}^{T(i)}}$, where $\mathbb{1}(\mathrm{a})$ is 1 if $a$ is true, and 0 otherwise. Intuitively, ${}_{r-1}\pi_{.j}^{T(i)}$ modifies ${}_{r-1}\pi_{kj}^{T(i)}$ to ensure that reads are only allocated to the transcripts in $C_j$.

II) Then, for $k = 1, \ldots, K$, we add each isoform counts across ECs to obtain the transcript level counts as: ${}_rX_k^{(i)} = \sum_{j=1}^J {}_rX_{kj}^{(i)}$, $k = 1, \ldots, K$ and $i = 1, \ldots, N$.

Update $\underline{\pi}|\delta, \underline{X}$: For $i = 1, \ldots, N$, we use the following Gibbs sampler:

$$ {}_r\pi^{(i)} | {}_{r-1}\delta, {}_rX^{(i)} \sim \mathcal{DIR}\left( {}_{r-1}\delta + {}_rX^{(i)} \right), \tag{S2} $$

where $\left( {}_{r-1}\delta + {}_rX^{(i)} \right) = \left( {}_{r-1}\delta_1 + {}_rX_1^{(i)}, \ldots, {}_{r-1}\delta_K + {}_rX_K^{(i)} \right)$.

Update $\delta|\underline{\pi}$: We draw our Metropolis proposal for $\delta$ as follows:

$$ log({}_r\delta) \sim \mathcal{N}\left( log({}_{r-1}\delta), {}_r\Sigma_\delta^{(prop)} \right), \tag{S3} $$

where ${}_r\Sigma_\delta^{(prop)}$ represents the ARW proposal matrix for $log(\delta)$ at the $r$-th iteration of the MCMC.

The proposed value $log({}_r\delta)$ is then accepted with probability:

$$ \frac{L_\delta({}_r\delta|{}_r\underline{\pi})\, f_N(log({}_r\delta)|\mu_\delta, \Sigma_\delta)}{L_\delta({}_{r-1}\delta|{}_r\underline{\pi})\, f_N(log({}_{r-1}\delta)|\mu_\delta, \Sigma_\delta)}, \tag{S4} $$

where $L_\delta(\delta|\underline{\pi}) = \prod_{i=1}^{N} f_{Dir}\left(\pi^{(i)}\middle|\,\delta\right)$, with $f_{Dir}(\cdot|\delta)$ being the density of the Dirichlet random variable with parameter $\delta$, and $f_N(\cdot|\mu_\delta, \Sigma_\delta)$ denotes the density of the multivariate normal distribution with mean vector $\mu_\delta$ and variance-covariance matrix $\Sigma_\delta$; $\mu_\delta$ and $\Sigma^2_\delta$ are determined according to whether an informative prior is formulated for the dispersion parameter, as explained in the Methods Section of the main manuscript.

The ARW matrix ${}_r\Sigma_\delta^{(prop)}$ is first updated after 200 iterations, and again when the *burn_in* is reached; in both cases the first 100 iterations are excluded from the covariance computation:

$$
{}_r\Sigma_\delta^{(prop)} = \begin{cases} diag(0.1, K) & \text{for } r \leq 200, \\ Cov(log({}_{101}\delta), \ldots, log({}_{200}\delta)) & \text{for } r \in \{201, \ldots, burn\_in\}, \\ Cov(log({}_{101}\delta), \ldots, log({}_{burn\_in}\delta)) & \text{for } r > burn\_in, \end{cases} \tag{S5}
$$

where $diag(a, b)$ represents the diagonal matrix of size $b$ with diagonal elements $a$, and $Cov(\cdot)$ indicates the variance-covariance matrix operator.

##### S1.2 EC with multiple genes

If an equivalence class has transcripts from multiple genes, we apply a minor change to the algorithm described in Section S1.1.

Updates of $\delta$ and $\underline{\pi}$ are still performed separately for every gene as shown in Section S1.1. In the sampling of $\underline{X}$, however, we modify ${}_{r-1}\pi_{.j}^{T(i)}$, in formula (S1), to include all transcripts from the genes in the $j$-th EC, with transcript level probabilities being weighted by the number of reads associated to each gene.

Assume the $j$-th EC has transcripts from two genes, $g_1$ and $g_2$, with $K_{g_1}$ and $K_{g_2}$ transcripts, respectively. At the $r$-th iteration of the MCMC, the probability vector ${}_{r-1}\pi_{.j}^{T(i)}$ in (S1) is replaced by:

$$
{}_{r-1}\tilde{\pi}_{.j}^{T(i)} = \left( {}_{r-1}\tilde{\pi}_{1jg_1}^{T(i)}, \ldots, {}_{r-1}\tilde{\pi}_{K_{g_1}jg_1}^{T(i)}, {}_{r-1}\tilde{\pi}_{1jg_2}^{T(i)}, \ldots, {}_{r-1}\tilde{\pi}_{K_{g_2}jg_2}^{T(i)} \right) \tag{S6}
$$

where the third subscript, $g_1$ or $g_2$, indicates the gene, and ${}_{r-1}\tilde{\pi}_{kjg}^{T(i)} = {}_{r-1}\pi_{kjg}^{T(i)} \sum_{k'=1}^{K_g} {}_rX_{k'g}^{(i)}$, for $k = 1, \ldots K_g$ and $g \in \{g_1, g_2\}$, with $\sum_{k'=1}^{K_g} {}_rX_{k'g}^{(i)}$ representing the total number of reads attributed to gene $g$ at the $r$-th iteration of the MCMC. The case with 3 or more genes is an EC is a natural extension of the one presented above.

##### S1.3 DTU test between 3 or more groups

When comparing 3 or more groups, parameters inference, which is performed separetely for each group, is identical to the case with 2 conditions, while DTU testing differs.

For simplicity consider the case with 3 groups, denoted by letters $A$, $B$ and $C$, with average transcript relative expression $\bar{\pi}_k^{TA}$, $\bar{\pi}_k^{TB}$ and $\bar{\pi}_k^{TC}$, respectively. For gene level testing, we consider the following system of hypothesis:

$$\begin{cases} \mathcal{H}_0: & \tilde{\omega}_k = 0, \text{ for } k \in \{1, \dots, 2K\} \\ \mathcal{H}_1: & \text{otherwise,} \end{cases} \tag{S7}$$

where $\tilde{\omega}_k = \bar{\pi}_k^{TG_1} - \bar{\pi}_k^{TG_2}$, for $k \in \{1, \dots, K\}$, and $\tilde{\omega}_k = \bar{\pi}_k^{TG_1} - \bar{\pi}_k^{TG_3}$, for $k \in \{K+1, \dots, 2K\}$, with $(G_1, G_2, G_3)$ being a permutation of the three groups $(A, B, C)$. In other words, to test if the average transcript proportions vary between groups, we choose a baseline group and compare the other two groups against it. The posterior distribution of $\tilde{\omega} = (\tilde{\omega}_1, \dots, \tilde{\omega}_{2K})$ can be approximated by a normal density [7], with mean $\hat{\tilde{\omega}}$ and variance matrix $\hat{\Sigma}_{\hat{\tilde{\omega}}}$, both inferred from the posterior chains. A multivariate Wald test [8] is implemented based on the null distribution of the test statistic: $\hat{\tilde{\omega}}_{-\{k',K+k'\}} \hat{\Sigma}^{-1}_{\hat{\tilde{\omega}}_{-\{k',K+k'\}}} \hat{\tilde{\omega}}^T_{-\{k',K+k'\}} \dot{\sim} \chi^2_{2K-2}$, where, as in the two-group comparison, $k' \in \{1, \dots, K\}$ is the transcript that should be removed from the test. Similarly, when individually testing the $k$-th transcript, we consider the system of hypothesis:

$$\begin{cases} \mathcal{H}_0: & \tilde{\omega}_k = 0, \text{ for } k \in \{k, K+k\} \\ \mathcal{H}_1: & \text{otherwise,} \end{cases} \tag{S8}$$

In this case we use a bivariate Wald test based on the statistic:
$\hat{\tilde{\omega}}_{\{k,K+k\}} \hat{\Sigma}^{-1}_{\hat{\tilde{\omega}}_{\{k,K+k\}}} \hat{\tilde{\omega}}^T_{\{k,K+k\}} \dot{\sim} \chi^2_2$. In both gene and transcript level tests, we alternatively use all 3 groups as baseline, with $(G_1, G_2, G_3) \in \{(A, B, C), (B, C, A), (C, A, B)\}$, and average the p-values of 3 tests.

DTU testing between more than 3 groups is a natural extension of the scenario illustrated above.

##### S1.4 Results details

The 3 vs. 3 simulated data is taken from Soneson et al. [9] were reads were simulated using Ensembl transcriptome version GRCh37.71; when simulating reads for the 6 vs. 6 simulation study we modified the pipeline in Soneson et al. [9] and kept the same Ensembl transcriptome version (GRCh37.71). Therefore, for the simulation studies, reads were then aligned using a filtered version of the GRCh37.71 genome and transcriptome: only transcripts with gene biotype equal to "protein_coding" and from "canonical" chromosomes (1 to 22, X and Y) were kept; furthermore duplicated transcripts (i.e., transcripts with exactly the same sequence) were removed.

Instead, for the experimental data analyses, we used the (unfiltered) Ensembl genome and transcriptome GRCh38.92, which was the latest version available when we ran the analyses.

For BANDITS, BayesDRIMSeq [10], DEXSeq_ECCs [11], DEXSeq_TECs [9], DRIMSeq [12] and rats [13], reads were first aligned via splice-aware genome aligner STAR [14], and then Salmon [15] was used on aligned reads to compute TECs and ECCs. For DEXSeq [16] and limma [17], reads were aligned via STAR, and then DEXSeq python function dexseq_prepare_annotation.py and dexseq_count.py were used to compute exon bin counts. For cjBitSeq [10, 18], reads were aligned with Bowtie2 [19].

BayesDRIMSeq and cjBitSeq scores represent decision rule $d_3$ in Papastamoulis and Rattray (2017) [10], and correspond to field *FDRraw* from the output files. The conservative scores BayesDRIMSeq_inv and cjBitSeq_inv indicate decision rule $d_4$ and refers to fields $fdrTrust$ and $FDR$, respectively, from the output files.

In the simulations study with transcript pre-filtering, we filtered isoforms based on Salmon TECs: we kept transcripts with least 10 counts (across all samples) and an average relative abundance of at least 0.01. The filtering was computed via BANDITS *filter_transcripts* function, with parameters *min_transcript_proportion = 0.01*, *min_transcript_counts = 10* and *min_gene_counts = 20*.

In gene-level plots, we excluded genes with less than 20 estimated counts across all samples. In transcript-level plots, we excluded transcripts with less than 10 estimated counts across all samples, and those belonging to a gene with less than 20 counts.

When stratifying results by gene expression, we computed the overall estimated abundance of each gene, across all samples, ranked them and split them into 3 equally sized groups. For the stratification in the 6 vs. 6 simulated data, we excluded genes with less than 1,200 estimated counts (i.e., 100 per sample on average), because no genes with less than 1,200 TECs are simulated to be differentially used.

In Figure 5 of the main manuscript, the blue component of panels A and C refers to the computational cost of STAR and Salmon (for BANDITS, BayesDRIMSeq, DEXSeq_ECCs, DEXSeq_TECs and DRIMSeq), or STAR and Salmon with 100 bootstrap replicates (for rats). For cjBitSeq, the blue component of panel A refers to the cost of Bowtie2, while in panel C it indicates the cost of STAR, Salmon (whose TECs are used to filter transcripts) and Bowtie2 on the filtered transcriptome. For DEXSeq and limma, the blue component of panel A refers to the cost of STAR and DEXSeq python functions (dexseq_prepare_annotation.py and dexseq_count.py), while in panel C it indicates the cost of STAR, Salmon (again, to filter transcripts), again STAR on the filtered transcriptome and DEXSeq python functions. rats is excluded from panels B and D because, although compatible with Salmon output, it requires bootstrap replicates.

##### S1.5 Visual inspection of two genes

To add biological perspective, we performed an in depth visual inspection of two genes from the “Best et al.” experimental data analysis. We selected two genes with adjusted p-value of 0.00 from BANDITS, one belonging to the set of 82 validated genes (ENSG00000147679) and one not previously validated (ENSG00000184432). Tables S9 and S10 report transcript-level adjusted p-values: BANDITS identifies two differentially used transcripts for gene ENSG00000184432 (ENST00000503326 and ENST00000507777) and three for gene ENSG00000147679 (ENST00000521071, ENST00000517820 and ENST00000521703).

Figures S13 and S16 illustrate the mean transcript-level proportions estimated from BANDITS, with 0.95 level profile Wald-type confidence intervals, while Figures S14, S15, S17 and S18 show sample-specific coverage and junction tracks obtained via IGV software [20]. BANDITS proportion plots show clear differences between groups in differentially used transcripts; some of these differences can also be easily visualized on the IGV plots, particularly those involving transcripts that are almost only expressed in one group.

These examples show how BANDITS can be effectively used to identify genes with differential transcript usage, as well as the individual transcripts that are affected.

### S2 Additional Tables and Figures

| Method | Variability between biological replicates | input data | ECs with >1 gene | mapping uncertainty modelled | Gene level test | Transcript level test | Transcript level proportions | Correct for transcript length | > 2 group comparisons | Allow for covariates |
|---|---|---|---|---|---|---|---|---|---|---|
| BANDITS | YES (DM) | ECCs | gene allocation sampled | YES (transcript allocation sampling) | YES | YES | YES | YES | YES | NO |
| BayesDRIMSeq [10] | YES (DM) | TECs | - | NO | YES | NO | NO | NO | NO | NO |
| cjBitSeq [10, 18] | NO (MN) | ECCs | counted once for each gene | YES (transcript allocation sampling) | YES | YES | YES | NO | NO | NO |
| DEXSeq [16] | YES (NB) | EBCs | - | - | YES | NO | NO | NO | YES | YES |
| DEXSeq on ECCs [11] | YES (NB) | ECCs | removed | - | YES | NO | NO | NO | YES | YES |
| DEXSeq on TECs [21] | YES (NB) | TECs | - | NO | YES | YES | NO | NO | YES | YES |
| DRIMSeq [12] | YES (DM) | TECs | - | NO | YES | YES | YES | NO | YES | YES |
| limma [17] | NO (LM) | EBCs | - | - | YES | NO | NO | NO | NO | YES |
| RATs [13] | NO (MN) | TECs | - | YES (bootstrap replicates of reads) | YES | YES | YES | NO | NO | NO |
| SUPPA2 [22] | YES | TECs | - | NO | YES | YES | YES | NO | NO | NO |

**Table S1** Main features of some of the most popular methods for DS, based on RNA-seq data. In the second column: DM = dirichlet-multinomial, MN = multinomial, NB = negative-binomial and LM = linear model. In the third column: ECCs = equivalence classes counts, TECs = transcript estimated counts, EBCs = exon bin counts. Note that "mapping uncertainty modelled" is missing in "DEXSeq", "limma" and "DEXSeq on ECs" rows, because inference is performed on EBCs and ECCs. Similarly, column "ECs with > 1 gene" is only applicable to methods working with equivalence classes (ECs). Note that "¿2 group comparison" excludes models, such as SUPPA2 and limma, that perform pairwise tests between all pairs of groups.

| tool | version |
|---|---|
| R | 3.6.0 |
| Bioconductor packages | 3.9 |
| Salmon | 0.9.0 |
| STAR | 2.5.1b |
| bowtie2 | 2.1.0 |
| cjBitSeq | 1.0 |
| BitSeq | 0.7.5 |
| SUPPA2 | 2.3 |
| RSEM | 1.2.21 |

**Table S2** Software versions used in all our analyses.

| | Low | | Mid | | High | |
|---|---|---|---|---|---|---|
| | Median position | AUC | Median position | AUC | Median position | AUC |
| BANDITS_inv | 374.00 | 0.79 | 178.00 | 0.78 | 172.00 | 0.88 |
| BANDITS | 430.50 | 0.77 | 262.00 | 0.76 | 168.00 | 0.87 |
| cjBitSeq | 339.50 | 0.82 | 348.00 | 0.74 | 241.00 | 0.86 |
| rats | 292.50 | 0.82 | 236.00 | 0.77 | 195.00 | 0.87 |
| DEXSeq_TECs | 507.50 | 0.78 | 352.00 | 0.74 | 156.00 | 0.88 |
| DEXSeq_ECCs | 288.50 | 0.75 | 421.00 | 0.74 | 201.00 | 0.87 |
| BayesDRIMSeq | 403.00 | 0.77 | 432.00 | 0.74 | 218.00 | 0.74 |
| DEXSeq | 336.50 | 0.79 | 684.00 | 0.76 | 255.00 | 0.81 |
| limma | 387.50 | 0.79 | 792.00 | 0.66 | 315.00 | 0.82 |
| SUPPA2 | 492.25 | 0.68 | 931.50 | 0.67 | 507.50 | 0.67 |
| DRIMSeq | 1637.75 | 0.55 | 2117.50 | 0.53 | 395.00 | 0.70 |
| cjBitSeq_inv | 1712.50 | 0.59 | 1717.00 | 0.58 | 1718.00 | 0.60 |
| BayesDRIMSeq_inv | 1826.50 | 0.53 | 1786.50 | 0.56 | 1750.00 | 0.61 |

**Table S3 Results from the "Best et al." experimental dataset, stratified by gene expression; methods are sorted by lowest "Median position" in the overall analysis (Table 1 of the main manuscript). "Median position" indicates the median position of the 83 validated genes in the ranking of 10,000 analyzed genes; AUC refers to the area under the ROC curve; pAUC 0.1 and 0.2 indicate the partial AUC of levels 0.1 and 0.2, respectively. Genes were separated in three equally sized groups according to their expression: "Low", "Mid" and "High".**

| | Gene test | | Transcript test | |
|---|---|---|---|---|
| | p-value | FDR | p-value | FDR |
| BANDITS | 2.17 | 0.22 | 5.85 | 0.16 |
| BANDITS_inv | 1.47 | 0.18 | - | - |
| BANDITS_maxGene | - | - | 0.18 | 0.05 |
| BayesDRIMSeq | 8.10 | 3.16 | - | - |
| BayesDRIMSeq_inv | - | 1.89 | - | - |
| cjBitSeq | 4.86 | 2.53 | 4.54 | 4.54 |
| cjBitSeq_inv | - | 0.92 | - | - |
| DEXSeq | - | 0.25 | - | - |
| DEXSeq_ECCs | - | 11.13 | - | - |
| DEXSeq_TECs | - | 3.37 | 9.75 | 1.09 |
| DRIMSeq | 3.61 | 0.35 | 4.25 | 0.21 |
| limma | 0.88 | 0.00 | - | - |
| rats | 50.41 | 49.61 | 54.60 | 50.60 |
| SUPPA2 | 13.14 | 1.40 | 4.34 | 0.44 |

**Table S4 Percentage of false positive tests returned from each method, at the 0.05 threshold, in the null experimental dataset.**

| | Low | Mid | High |
|---|---|---|---|
| BANDITS | 0.05 | 0.19 | 0.42 |
| BANDITS_inv | 0.05 | 0.16 | 0.33 |
| BayesDRIMSeq | 1.46 | 6.71 | 1.30 |
| BayesDRIMSeq_inv | 1.09 | 3.93 | 0.63 |
| cjBitSeq | 0.20 | 2.57 | 4.81 |
| cjBitSeq_inv | 0.10 | 1.01 | 1.67 |
| DEXSeq | 0.01 | 0.04 | 0.70 |
| DEXSeq_ECCs | 0.25 | 3.42 | 29.73 |
| DEXSeq_TECs | 0.74 | 3.61 | 5.78 |
| DRIMSeq | 0.16 | 0.42 | 0.45 |
| limma | 0.00 | 0.00 | 0.00 |
| rats | 8.73 | 57.44 | 82.65 |
| SUPPA2 | 0.75 | 1.65 | 1.79 |

**Table S5 Percentage of false positive tests returned from each method in the null experimental dataset, stratified by gene expression, according to gene-level FDR with a 0.05 threshold. Genes were separated in three equally sized groups according to their expression: "Low", "Mid" and "High".**

| | Unfiltered | Filtered |
|---|---|---|
| Salmon | 42 | - |
| Salmon_boot | 148 | - |
| STAR | 271 | 254 |
| BANDITS | 174 | 59 |
| BayesDRIMSeq | 29 | 18 |
| DEXSeq_ECCs | 43 | 39 |
| DEXSeq_TECs | 12 | 3 |
| DRIMSeq | 13 | 10 |
| rats | 29 | 27 |
| bowtie2 | 1111 | 811 |
| cjBitSeq | 4896 | 3401 |
| DEXSeq python | 1871 | 1740 |
| DEXSeq R | 61 | 30 |
| limma | 2 | 1 |

**Table S6 Computational cost, expressed in minutes, of each individual step. Columns "Unfiltered" and "Filtered" refer to the analyses run on the original transcriptome and on the filtered one, respectively. "Salmon" and "Salmon_boot" refer to running Salmon on the transcript alignments computed from STAR; "Salmon_boot" additionally computes 100 bootstrap replicates (used by rats). "DEXSeq python" indicates the python functions dexseq_prepare_annotation.py and dexseq_count.py, while "DEXSeq R" refers to the pipeline of DEXSeq computed in R.**

| | Unfiltered | Filtered |
|---|---|---|
| cjBitSeq | 6007 | 4524 |
| DEXSeq | 2203 | 2337 |
| limma | 2143 | 2308 |
| rats | 448 | 446 |
| BANDITS | 487 | 371 |
| DEXSeq_ECCs | 355 | 352 |
| BayesDRIMSeq | 341 | 330 |
| DRIMSeq | 326 | 322 |
| DEXSeq_TECs | 325 | 316 |

**Table S7 Overall computational cost, expressed in minutes, of the full pipeline of each method, including alignment and quantification steps. Columns "Unfiltered" and "Filtered" refer to the analyses run on the original transcriptome and on the filtered one, respectively. Methods are sorted by "Filtered" times.**

| | Unfiltered | Filtered |
|---|---|---|
| Salmon | 2.4 | 2.4 |
| Salmon_boot | 3.1 | 3.1 |
| STAR | 34.7 | 33.6 |
| BANDITS | 1.8 | 0.9 |
| BayesDRIMSeq | 0.7 | 0.7 |
| DEXSeq_ECCs | 4.7 | 4.2 |
| DEXSeq_TECs | 2.0 | 1.5 |
| DRIMSeq | 0.7 | 0.7 |
| rats | 5.8 | 3.3 |
| bowtie2 | 0.8 | 0.6 |
| cjBitSeq | 3.4 | 2.2 |
| DEXSeq python | 0.8 | 0.4 |
| DEXSeq R | 10.2 | 5.2 |
| limma | 1.4 | 1.2 |

**Table S8 Maximum RAM, expressed in gigabytes, used in each individual step. Columns "Unfiltered" and "Filtered" refer to the analyses run on the original transcriptome and on the filtered one, respectively. "Salmon" and "Salmon_boot" refer to running Salmon on the transcript alignments computed from STAR; "Salmon_boot" additionally computes 100 bootstrap replicates (used by rats). "DEXSeq python" indicates the python functions dexseq_prepare_annotation.py and dexseq_count.py, while "DEXSeq R" refers to the pipeline of DEXSeq computed in R.**

| Transcript_ID | Adjusted p-value |
|---|---|
| ENST00000503326 | 0.00 |
| ENST00000507777 | 0.00 |
| ENST00000333188 | 0.05 |
| ENST00000514508 | 1.00 |
| ENST00000512242 | 1.00 |
| ENST00000510181 | 1.00 |
| ENST00000510491 | 1.00 |
| ENST00000513274 | 1.00 |
| ENST00000502734 | 1.00 |
| ENST00000504295 | 1.00 |
| ENST00000512153 | 1.00 |
| ENST00000515006 | 1.00 |
| ENST00000512309 | 1.00 |

**Table S9 Transcript-level adjusted p-values obtained from BANDITS for gene ENSG00000184432. Gene-level adjusted p-value is 0.00.**

| Transcript_ID | Adjusted p-value |
|---|---|
| ENST00000521071 | 0.00 |
| ENST00000517820 | 0.00 |
| ENST00000521703 | 0.00 |
| ENST00000309822 | 0.18 |
| ENST00000521974 | 0.63 |
| ENST00000520733 | 1.00 |
| ENST00000519443 | 1.00 |
| ENST00000524128 | 1.00 |
| ENST00000517814 | 1.00 |

**Table S10 Transcript-level adjusted p-values obtained from BANDITS for gene ENSG00000147679. Gene-level adjusted p-value is 0.00.**

| | Median position | AUC | pAUC 0.1 | pAUC 0.2 | top 100 | top 200 | GO 0.01 | GO 0.05 |
|---|---|---|---|---|---|---|---|---|
| BANDITS | 673 | 0.80 | 0.04 | 0.11 | 18 | 25 | 0.34 | 0.33 |
| BANDITS_inv | 596 | 0.81 | 0.04 | 0.11 | 16 | 24 | 0.32 | 0.35 |
| BANDITS_NoPrior | 759 | 0.79 | 0.04 | 0.10 | 17 | 24 | 0.33 | 0.30 |
| BANDITS_NoPrior_inv | 672 | 0.80 | 0.04 | 0.11 | 17 | 24 | 0.30 | 0.33 |

**Table S11 Results from the "Best et al." experimental dataset. "BANDITS_NoPrior" refers to BANDITS being run with vaguely-informative prior (default when no informative prior is provided). "Median position" indicates the median position of the 83 validated genes in the ranking of 10,000 analyzed genes; AUC refers to the area under the ROC curve; pAUC 0.1 and 0.2 represent the partial AUC of levels 0.1 and 0.2, respectively; "top 100" and "top 200" report the number of validated genes (82 in total) in the 100 and 200 genes with lowest FDR from each method; "GO 0.01" and "GO 0.05" indicate the fraction of "validated GO terms" found by each method, when considering FDR thresholds 0.01 and 0.05, respectively.**

| | Gene test | | Transcript test | |
|---|---|---|---|---|
| | p-value | FDR | p-value | FDR |
| BANDITS | 2.17 | 0.22 | 5.85 | 0.16 |
| BANDITS_inv | 1.47 | 0.18 | - | - |
| BANDITS_maxGene | - | - | 0.18 | 0.05 |
| BANDITS_NoPrior | 6.12 | 1.57 | 9.38 | 0.82 |
| BANDITS_NoPrior_inv | 3.46 | 1.20 | - | - |
| BANDITS_NoPrior_maxGene | - | - | 0.31 | 0.43 |

**Table S12 Percentage of false positive tests returned from each method, at the 0.05 threshold, in the null experimental dataset. "BANDITS_NoPrior" refers to BANDITS being run with vaguely-informative prior (default when no informative prior is provided).**

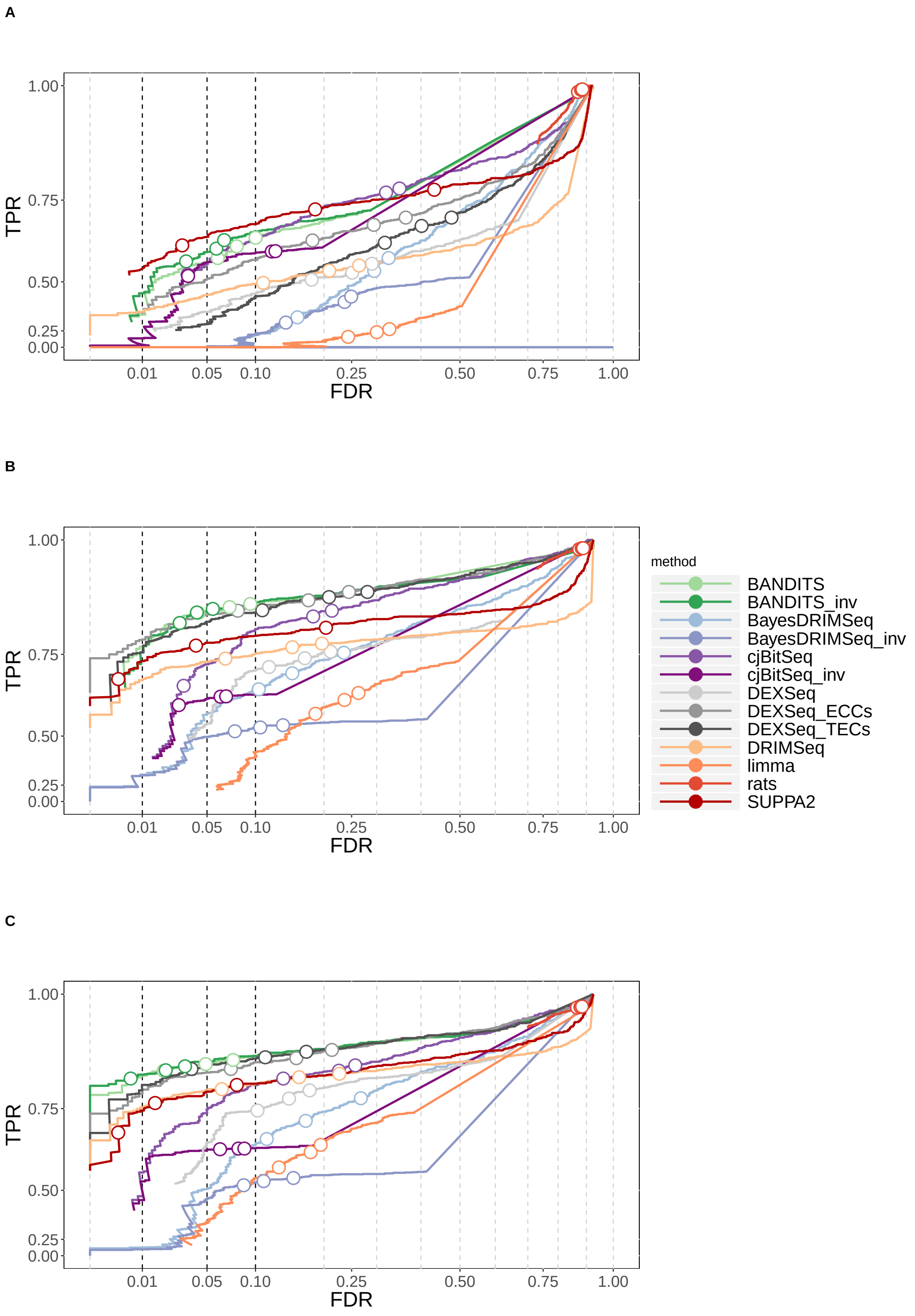

**Figure S1** TPR vs. FDR for gene-level testing for the three 2-group comparison simulation studies. A) 3 vs. 3 simulation study; B) 6 vs. 6 simulation study; C) 6 vs. 6 simulation study with transcript pre-filtering (transcripts with at least 10 counts and an average relative abundance of 0.01). Circles indicate observed FDR for 0.01, 0.05 and 0.1 significance thresholds.

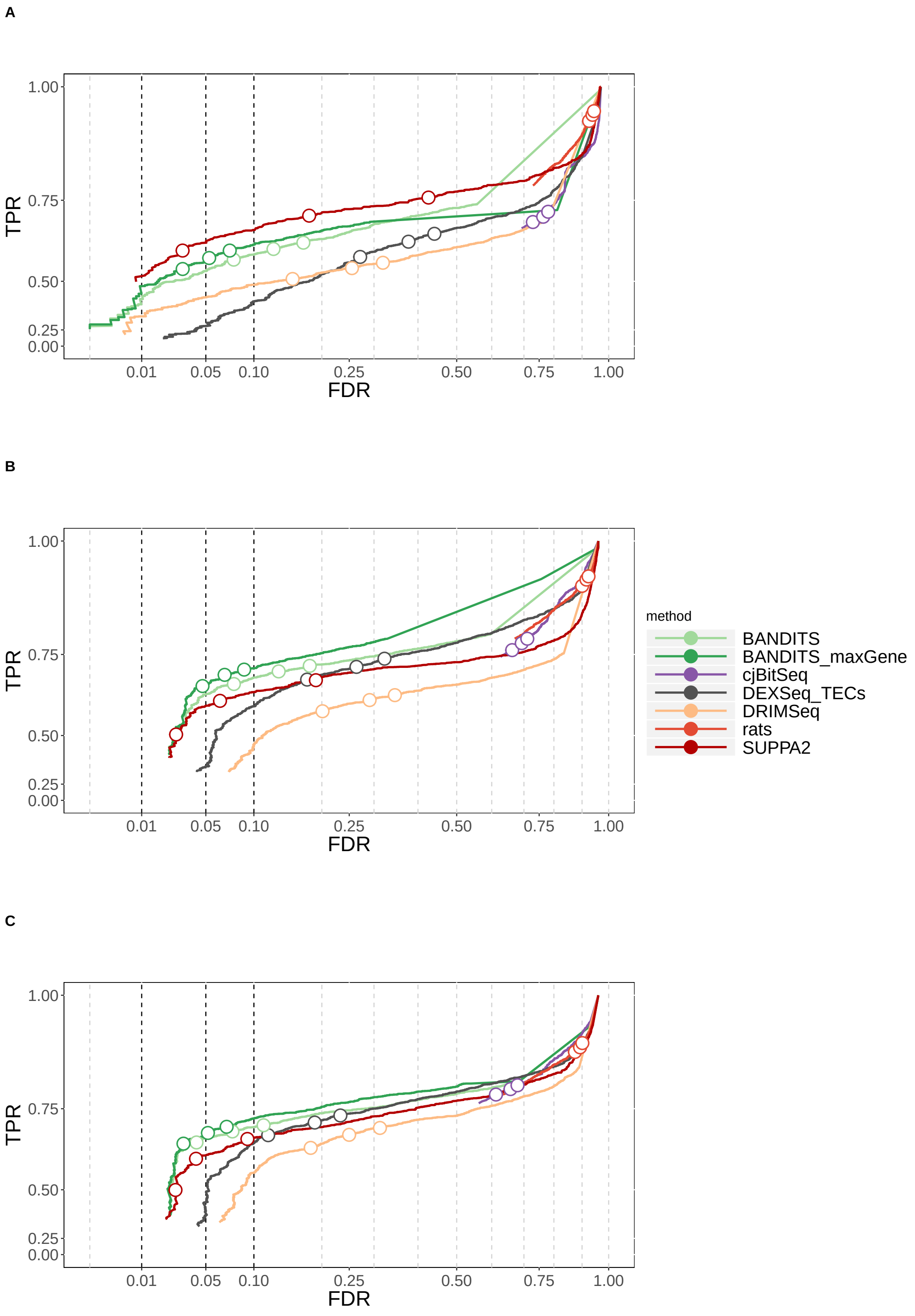

**Figure S2** TPR vs. FDR for transcript-level testing for the three 2-group comparison simulation studies. A) 3 vs. 3 simulation study; B) 6 vs. 6 simulation study; C) 6 vs. 6 simulation study with transcript pre-filtering (transcripts with at least 10 counts and an average relative abundance of 0.01). Circles indicate observed FDR for 0.01, 0.05 and 0.1 significance thresholds. Note that, for cjBitSeq, we considered the probability that a transcript is not differentially used, which does not guarantee FDR control.

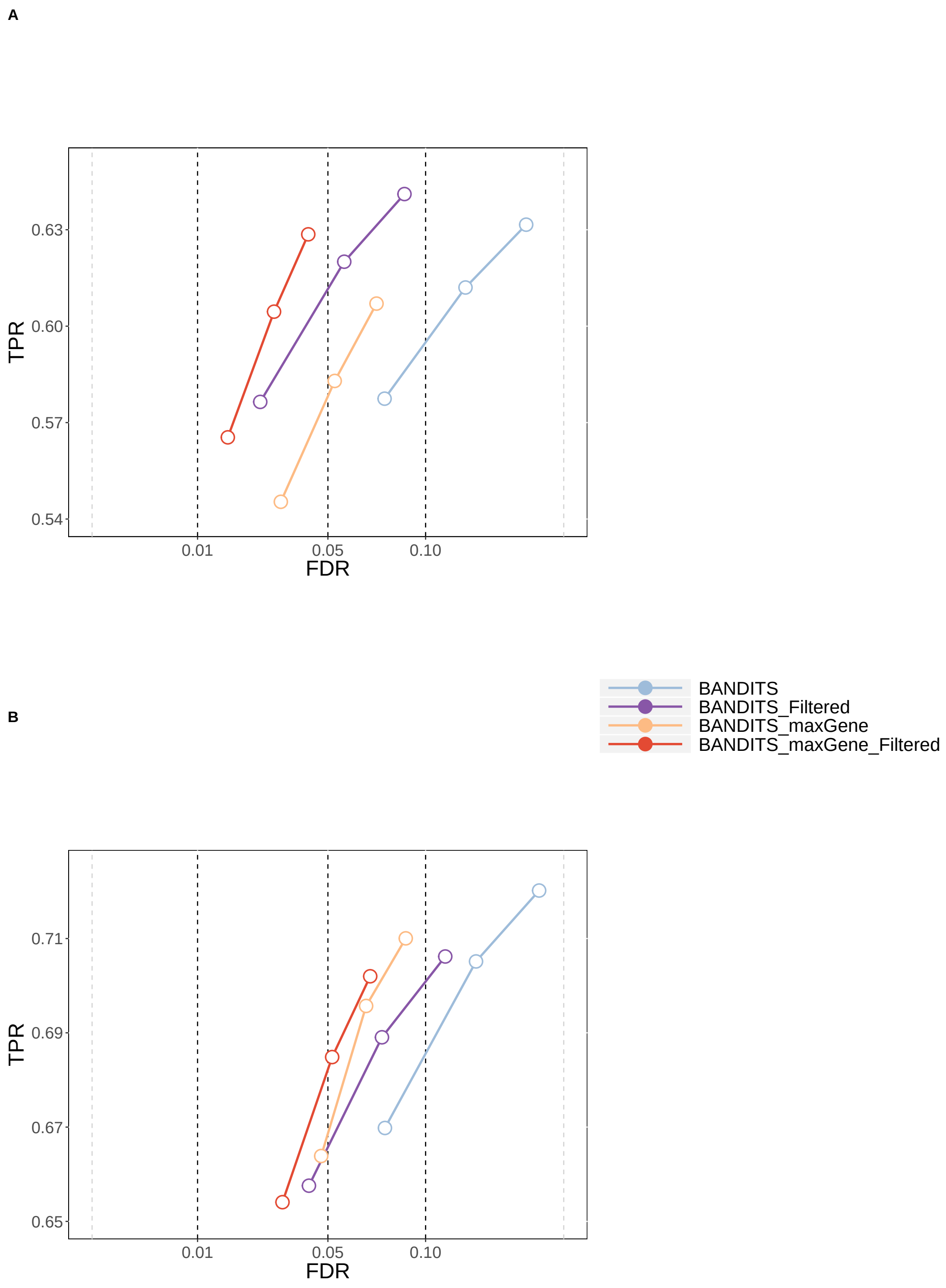

**Figure S3** TPR vs. FDR for gene-level testing, showing the improvement in performance of BANDITS results when pre-filtering transcripts. A) 3 vs. 3 simulation study; B) 6 vs. 6 simulation study. Circles indicate observed FDR for 0.01, 0.05 and 0.1 significance thresholds.

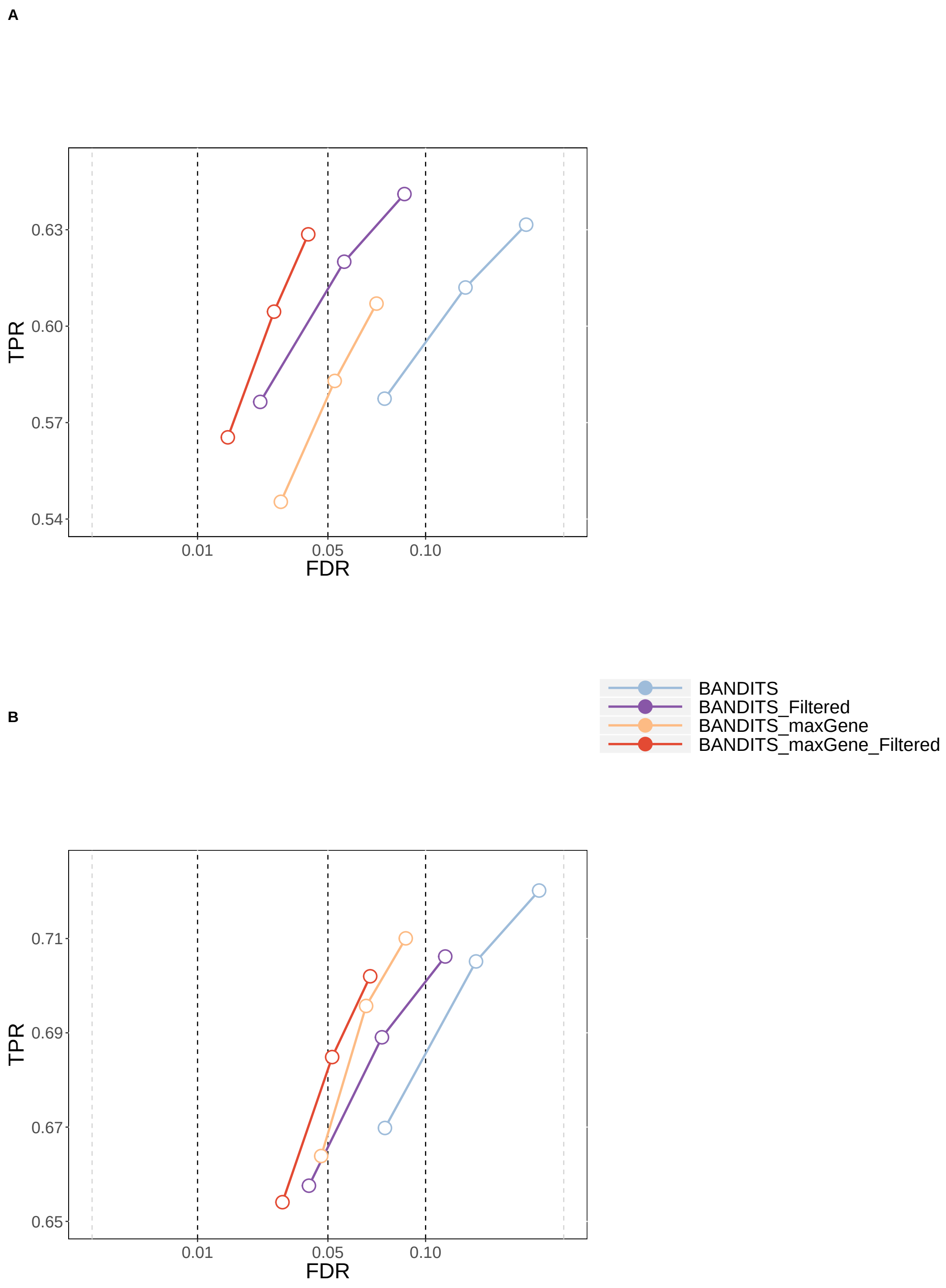

**Figure S4** TPR vs. FDR for transcript-level testing, showing the improvement in performance of BANDITS results when pre-filtering transcripts. A) 3 vs. 3 simulation study; B) 6 vs. 6 simulation study. Circles indicate observed FDR for 0.01, 0.05 and 0.1 significance thresholds.

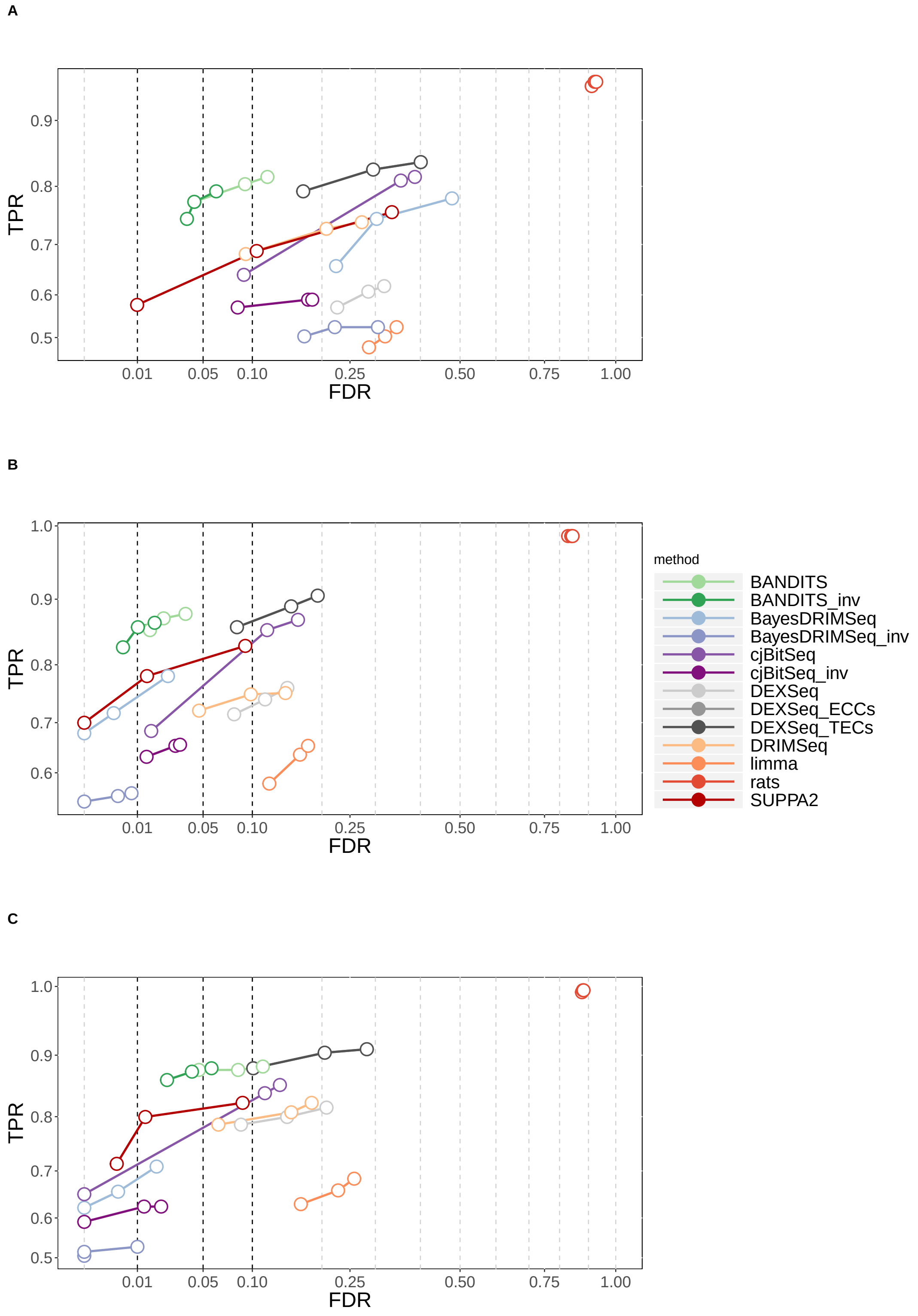

**Figure S5** TPR vs. FDR for gene-level testing in the 6 vs. 6 simulation study, stratified by gene expression. A) Low expression; B) medium expression; C) high expression. Circles indicate observed FDR for 0.01, 0.05 and 0.1 significance thresholds.

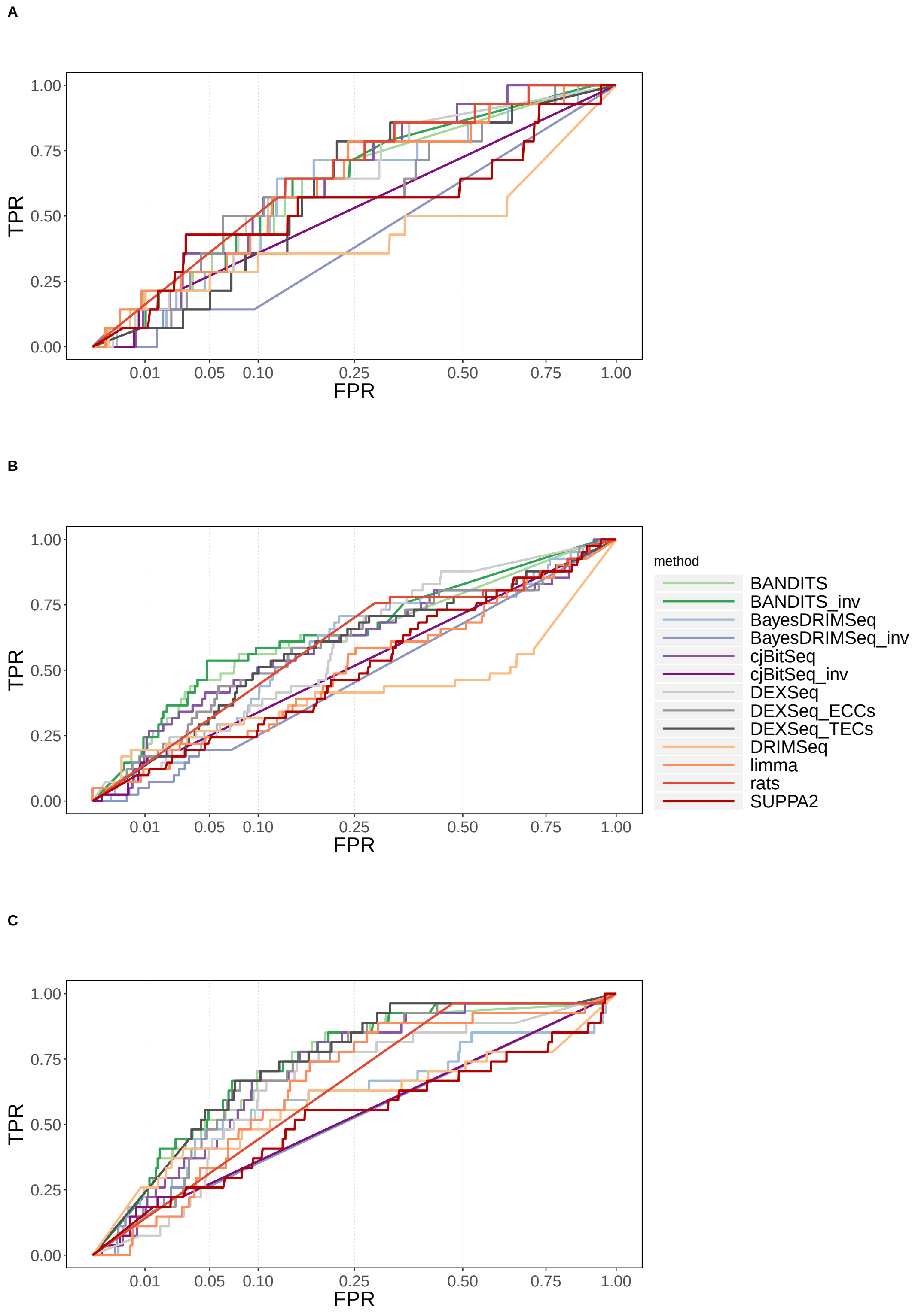

**Figure S6** ROC curve (TPR vs. FPR) for gene-level testing in the "Best et al." experimental dataset, stratified by gene expression. A) Low expression; B) medium expression; C) high expression.

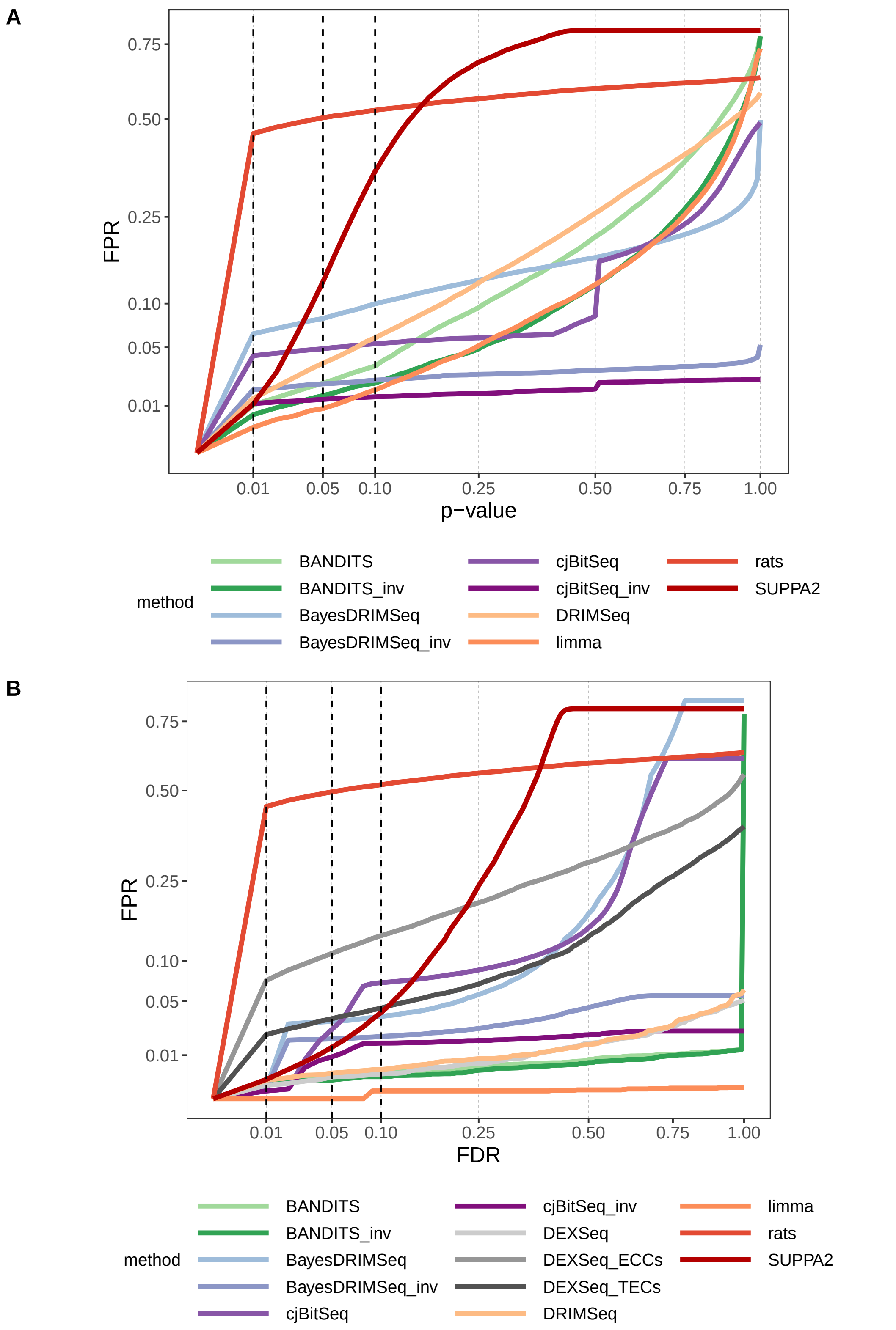

**Figure S7** FPRs for gene-level testing in the null experimental dataset. A) FPR vs. p-value; B) FPR vs. FDR. In panel A), we considered the minimum of transcript-level raw p-values to obtain a SUPPA2 gene-level p-value.

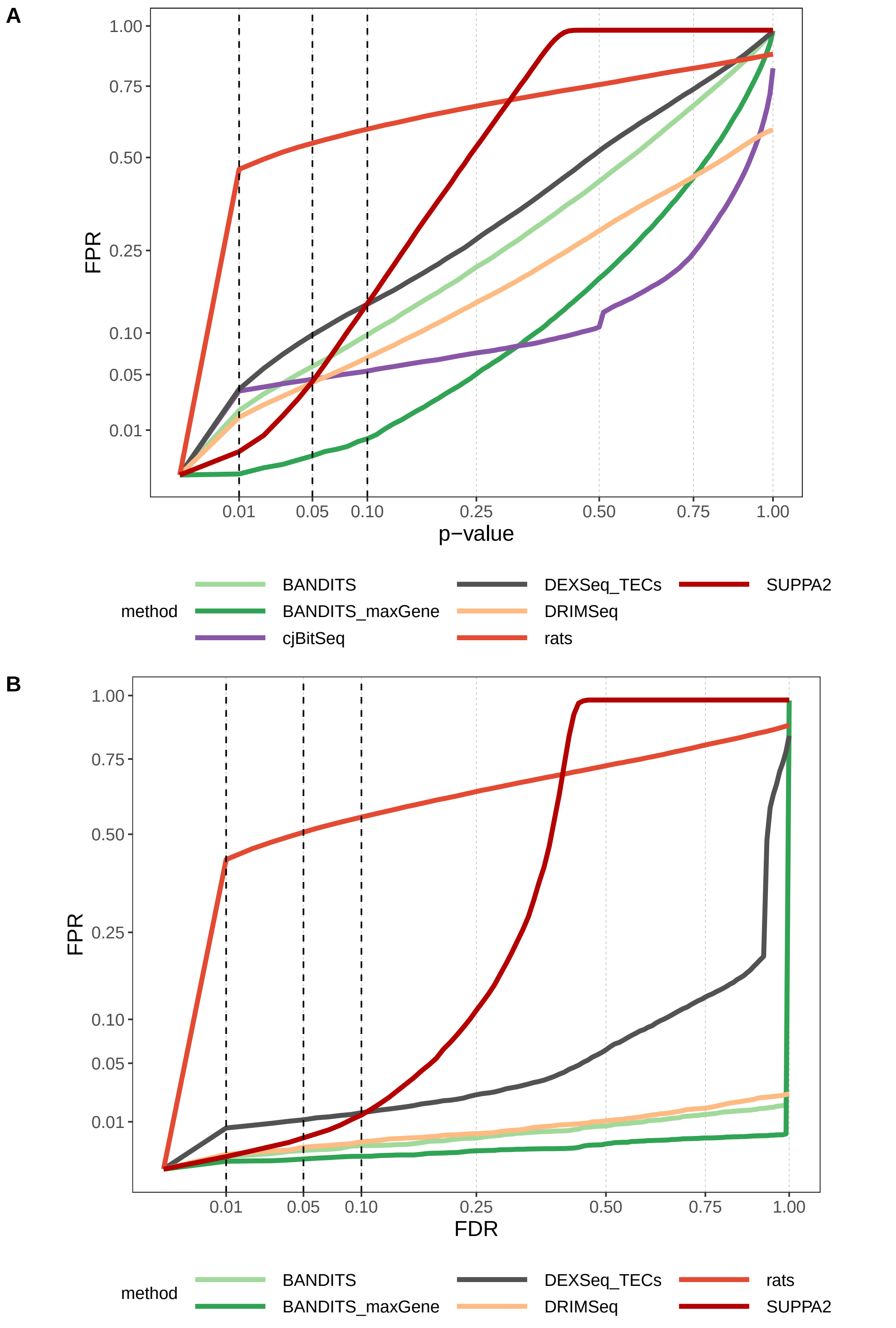

**Figure S8** FPRs for transcript-level testing in the null experimental dataset. A) FPR vs. p-value; B) FPR vs. FDR Note that, for cjBitSeq, in both panels, we considered the probability that a transcript is not differentially used, which does not guarantee FDR control.

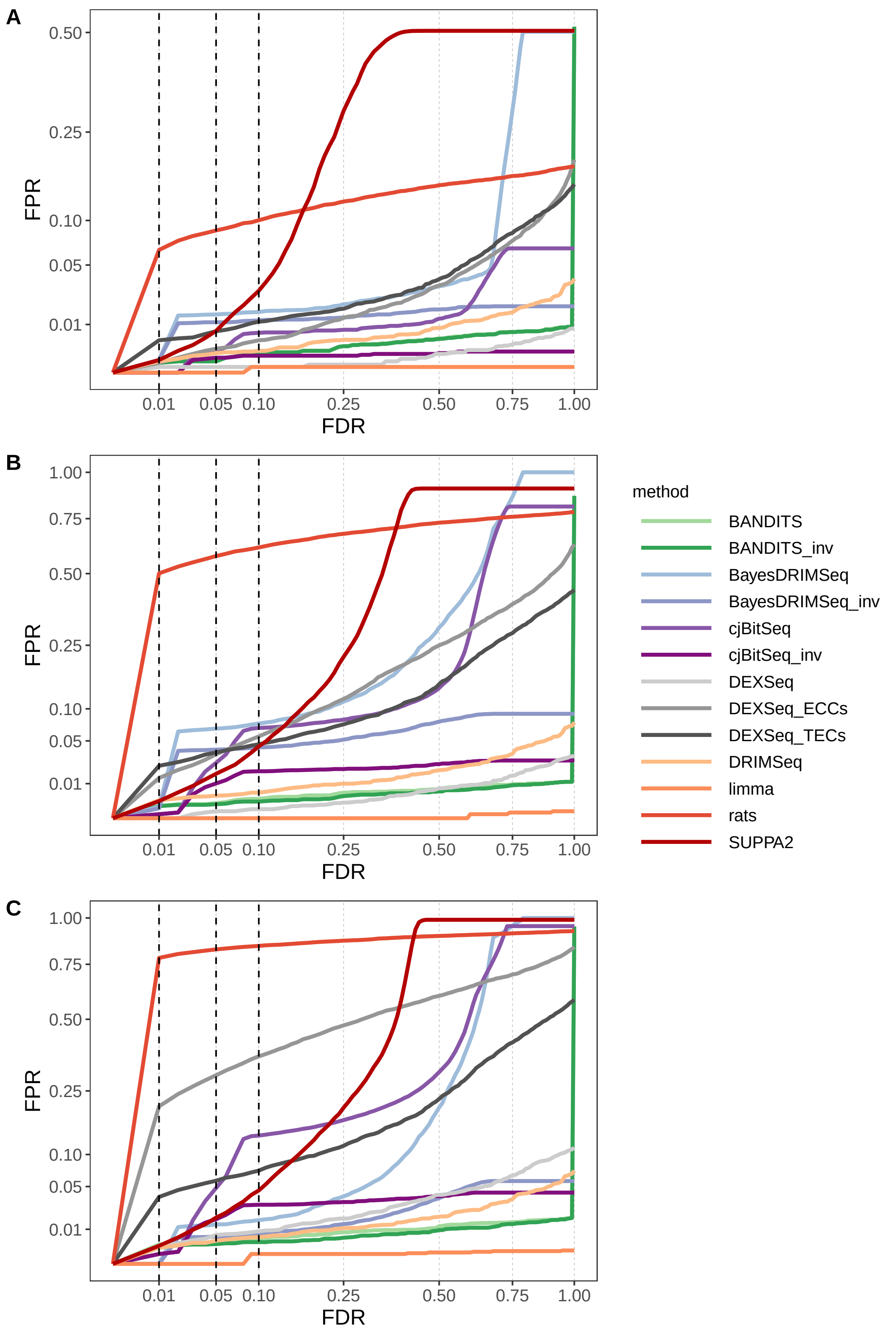

**Figure S9** FPR vs. FDR for gene-level testing in the null experimental dataset, stratified by gene expression. A) Low expression; B) medium expression; C) high expression.

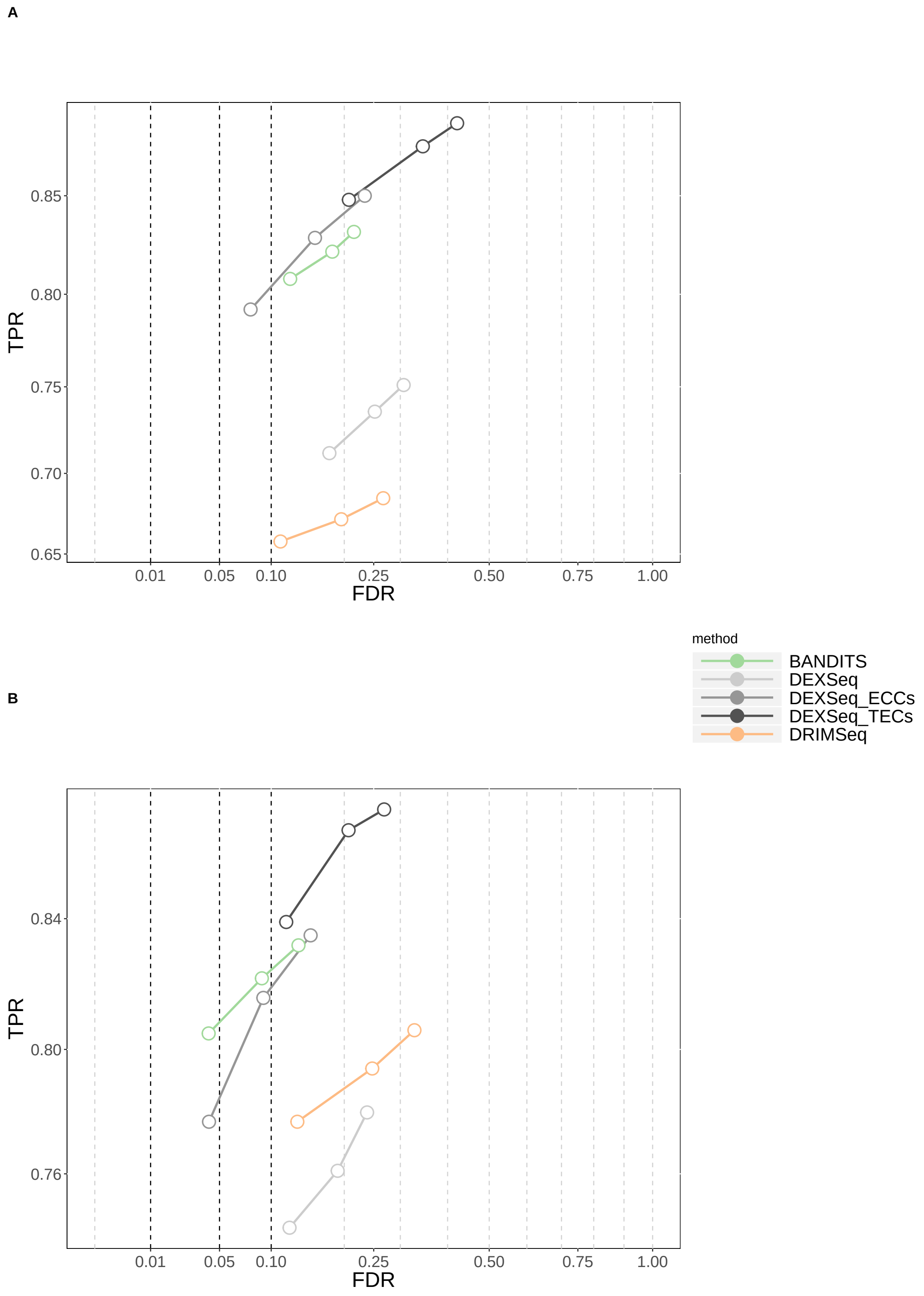

**Figure S10** TPR vs. FDR for gene-level testing for the 3-group comparison simulation study. A) 3 vs. 3 vs. 6 simulation study; B) 3 vs. 3 vs. 6 simulation study with transcript pre-filtering (transcripts with at least 10 counts and an average relative abundance of 0.01). Circles indicate observed FDR for 0.01, 0.05 and 0.1 significance thresholds.

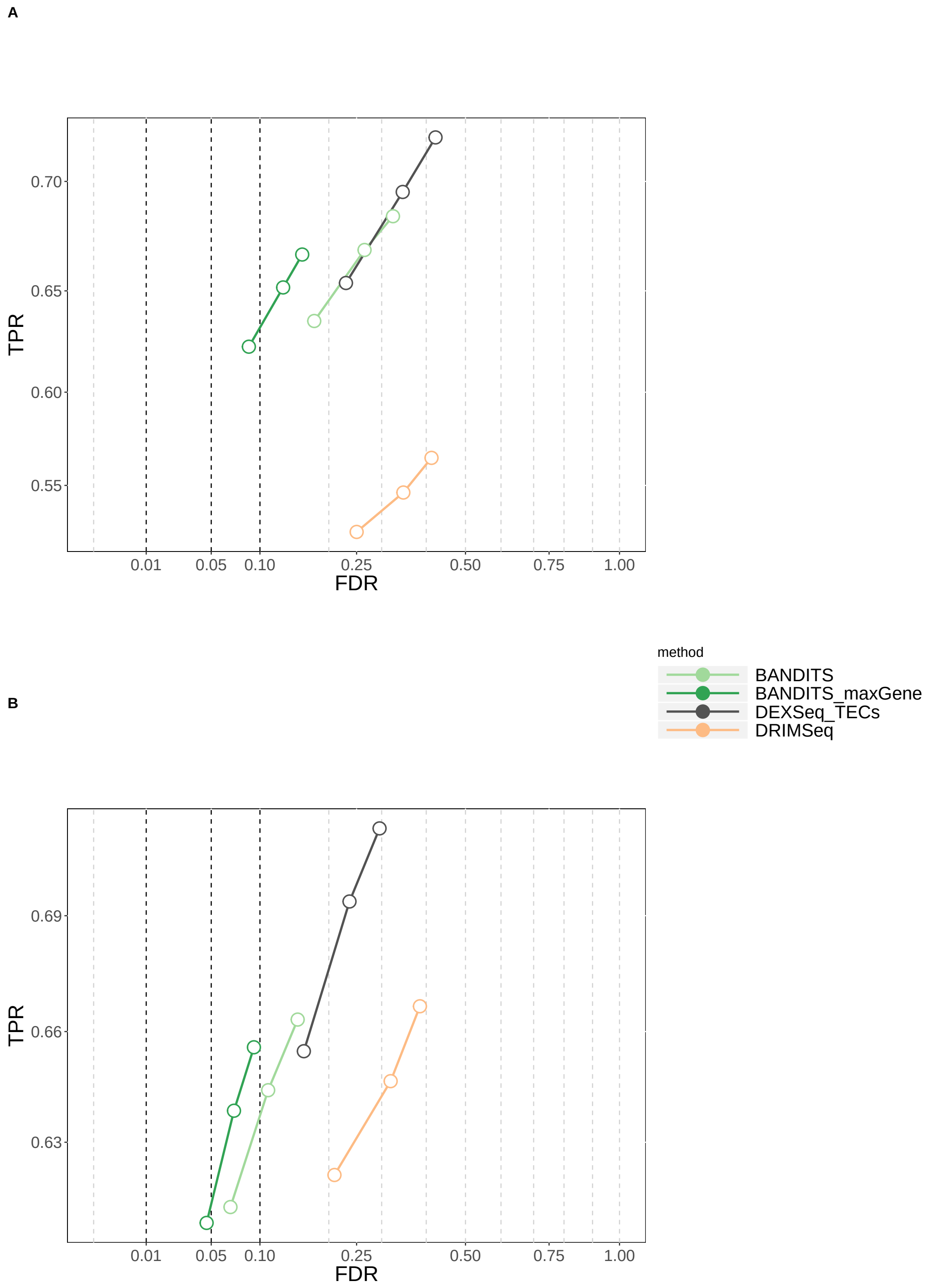

**Figure S11** TPR vs. FDR for transcript-level testing for the 3-group comparison simulation study. A) 3 vs. 3 vs. 6 simulation study; B) 3 vs. 3 vs. 6 simulation study with transcript pre-filtering (transcripts with at least 10 counts and an average relative abundance of 0.01). Circles indicate observed FDR for 0.01, 0.05 and 0.1 significance thresholds.

A

B

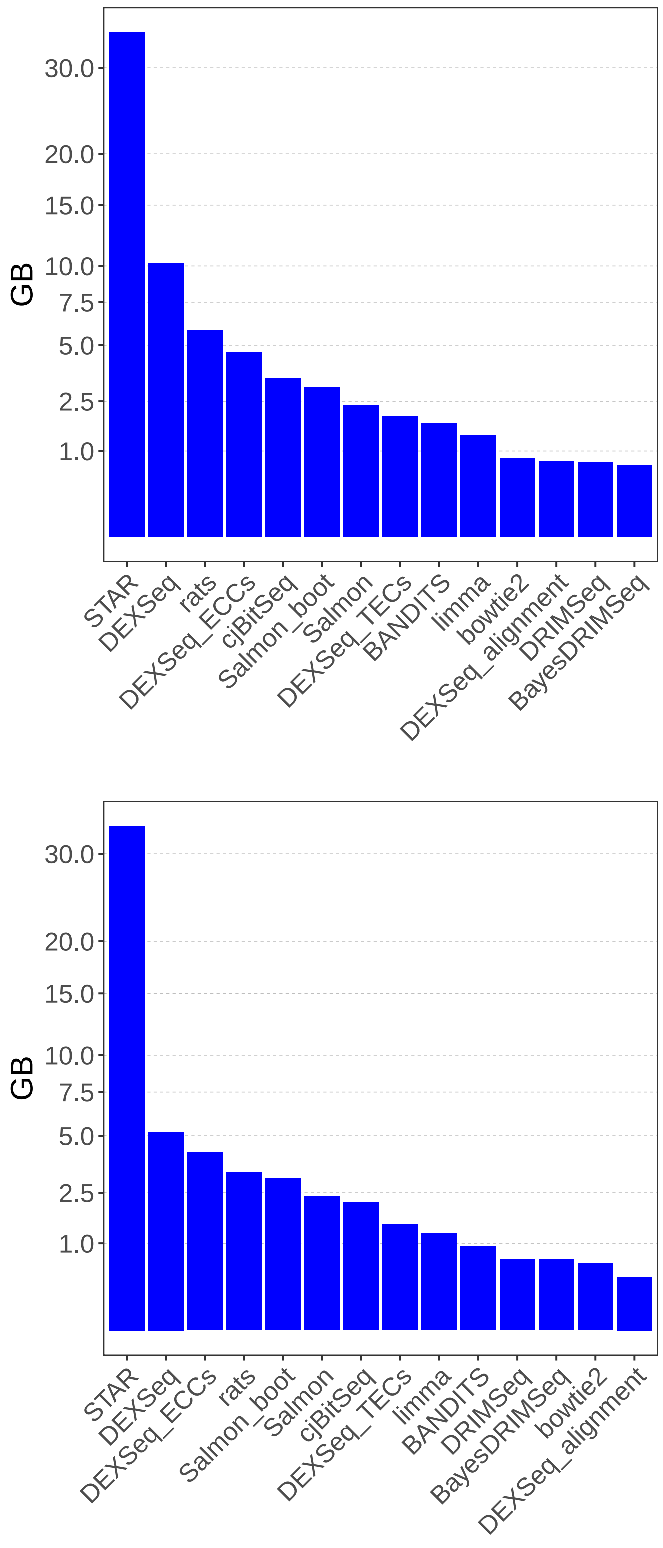

**Figure S12** Maximum RAM, expressed in gigabytes, used in each individual step. A) 6 vs. 6 simulation study; B) 6 vs. 6 simulation study with transcript pre-filtering (transcripts with at least 10 counts and an average relative abundance of 0.01). "Salmon" and "Salmon_boot" refer to running Salmon on the transcript alignments computed from STAR; "Salmon_boot" additionally computes 100 bootstrap replicates (used by rats).

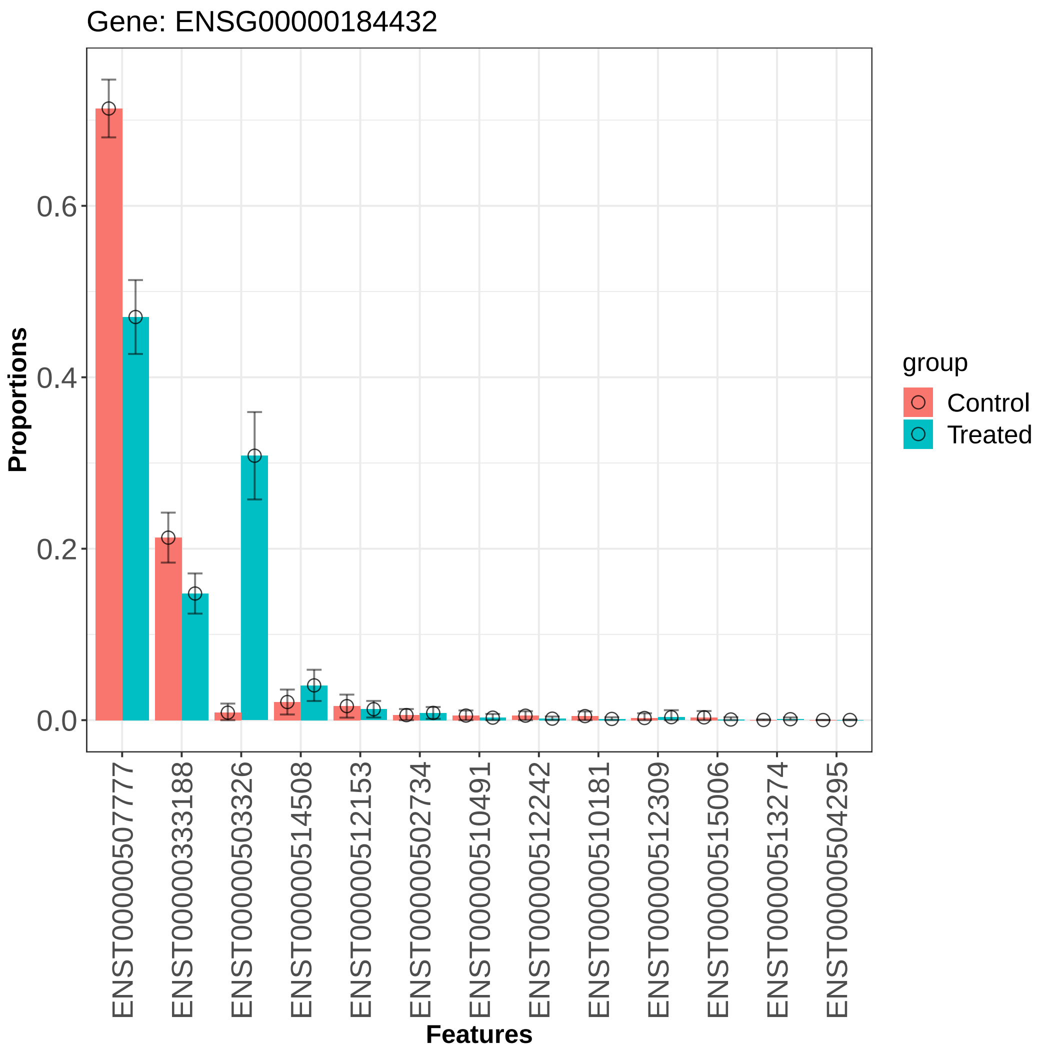

**Figure S13** Average probability of expressing transcripts, $\bar{\pi}^T$, for groups "Controls" and "Treated", estimated from the "Best et al." experimental data with BANDITS for gene ENSG00000184432. The vertical bars indicate 0.95 level profile Wald type confidence intervals. Image realized via *plot_proportions* function from BANDITS Bioconductor package. Gene ENSG00000184432 and transcripts ENST00000503326 and ENST00000507777 are identified as differentially used between conditions (adjusted p-value below 0.0001 in all three cases). No other transcripts is detected as significant (adjusted p-value above 0.05).

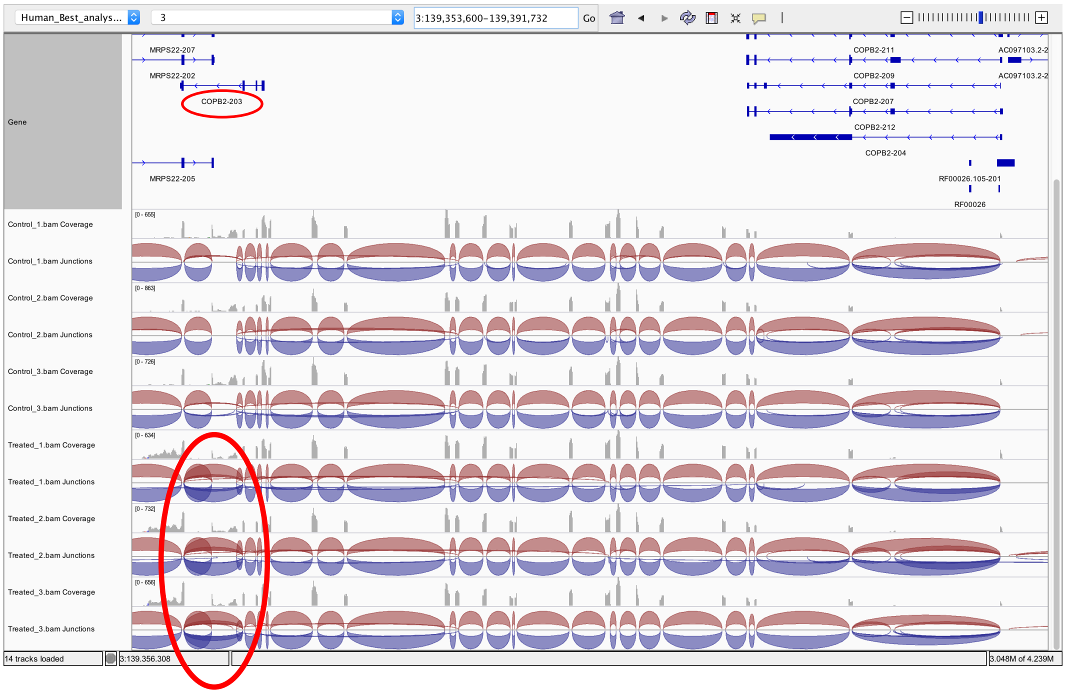

**Figure S14** IGV browser track visualization of coverage and junction reads for the entire gene ENSG00000184432. The junction tracks on the left area of the plot (inside the red circle) show that significantly more reads from Treated samples are compatible with transcript ENST00000503326 (COPB2-203, annotation in the top left area of the image), indicating differential usage of this transcript.

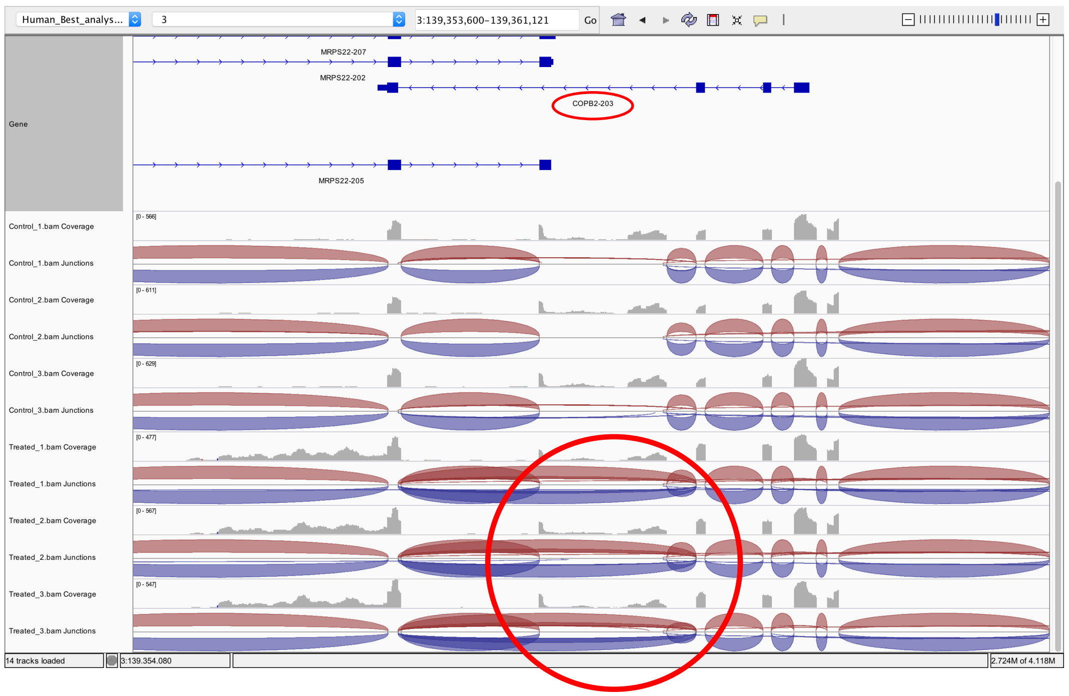

**Figure S15** IGV browser track visualization of coverage and junction reads for gene ENSG00000184432, zommed in the region around differentially used transcript ENST00000503326 (COPB2-203). The junction tracks on the central area of the plot (inside the red circle) show that significantly more reads from Treated samples are compatible with transcript ENST00000503326 (COPB2-203, annotation in the top central area of the image), indicating differential usage of this transcript.

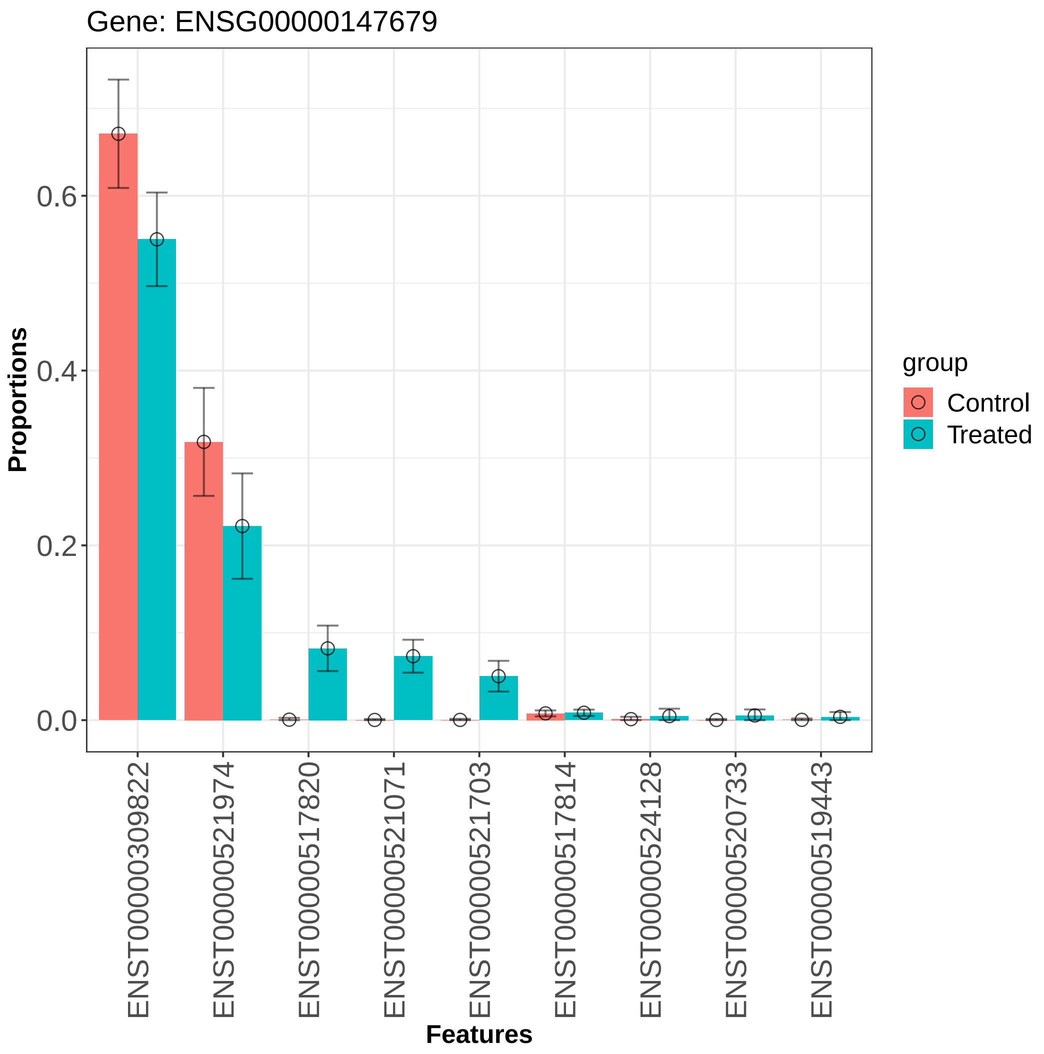

**Figure S16** Average probability of expressing transcripts, $\bar{\pi}^T$, for groups "Controls" and "Treated", estimated from the "Best et al." experimental data with BANDITS for gene ENSG00000147679. The vertical bars indicate 0.95 level profile Wald type confidence intervals. Image realized via *plot_proportions* function from BANDITS Bioconductor package. Gene ENSG00000147679 and transcripts ENST00000521071, ENST00000517820 and ENST00000521703 are identified as differentially used between conditions (adjusted p-value below 0.0001 in all four cases). No other transcripts is detected as significant (adjusted p-value above 0.05).

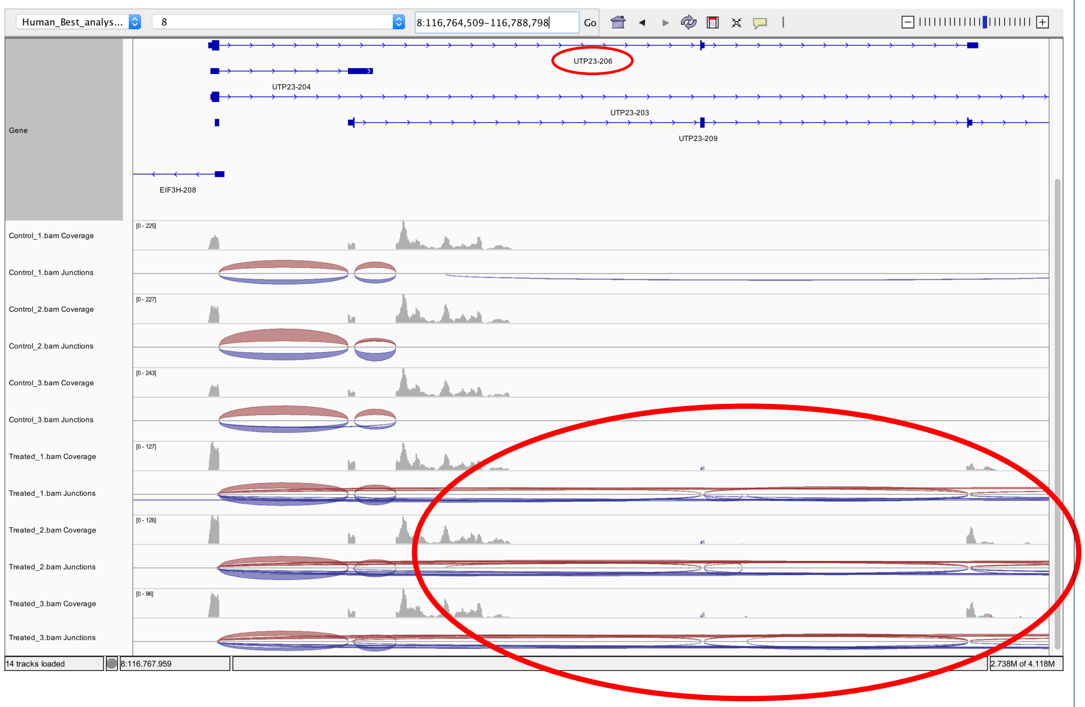

**Figure S17** IGV browser track visualization of coverage and junction reads for gene ENSG00000147679, zommed in the region around differentially used transcript ENST00000521071 (UTP23-206). The exon coverage and junction tracks on the right area of the plot (inside the red circle) clearly indicate that Treated samples undergo differential usage. The transcripts involved, not all visible due to their lengths, are ENST00000521071 (UTP23-206), ENST00000517820 (UTP23-203) and ENST00000521703 (UTP23-207).

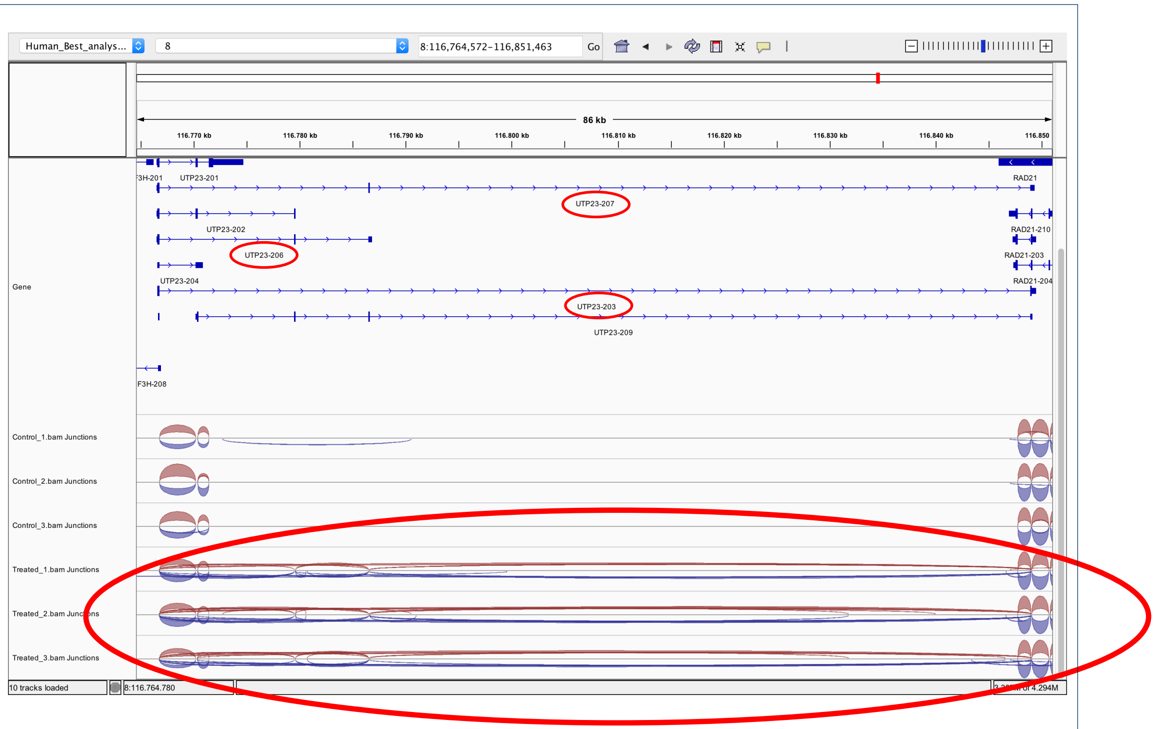

**Figure S18** IGV browser track visualization of junction reads for gene ENSG00000147679, zommed in the region around differentially used transcripts ENST00000521071 (UTP23-206), ENST00000517820 (UTP23-203) and ENST00000521703 (UTP23-207). The exon junction tracks on the bottom area of the plot (inside the red circle) clearly indicate that Treated samples undergo differential usage.

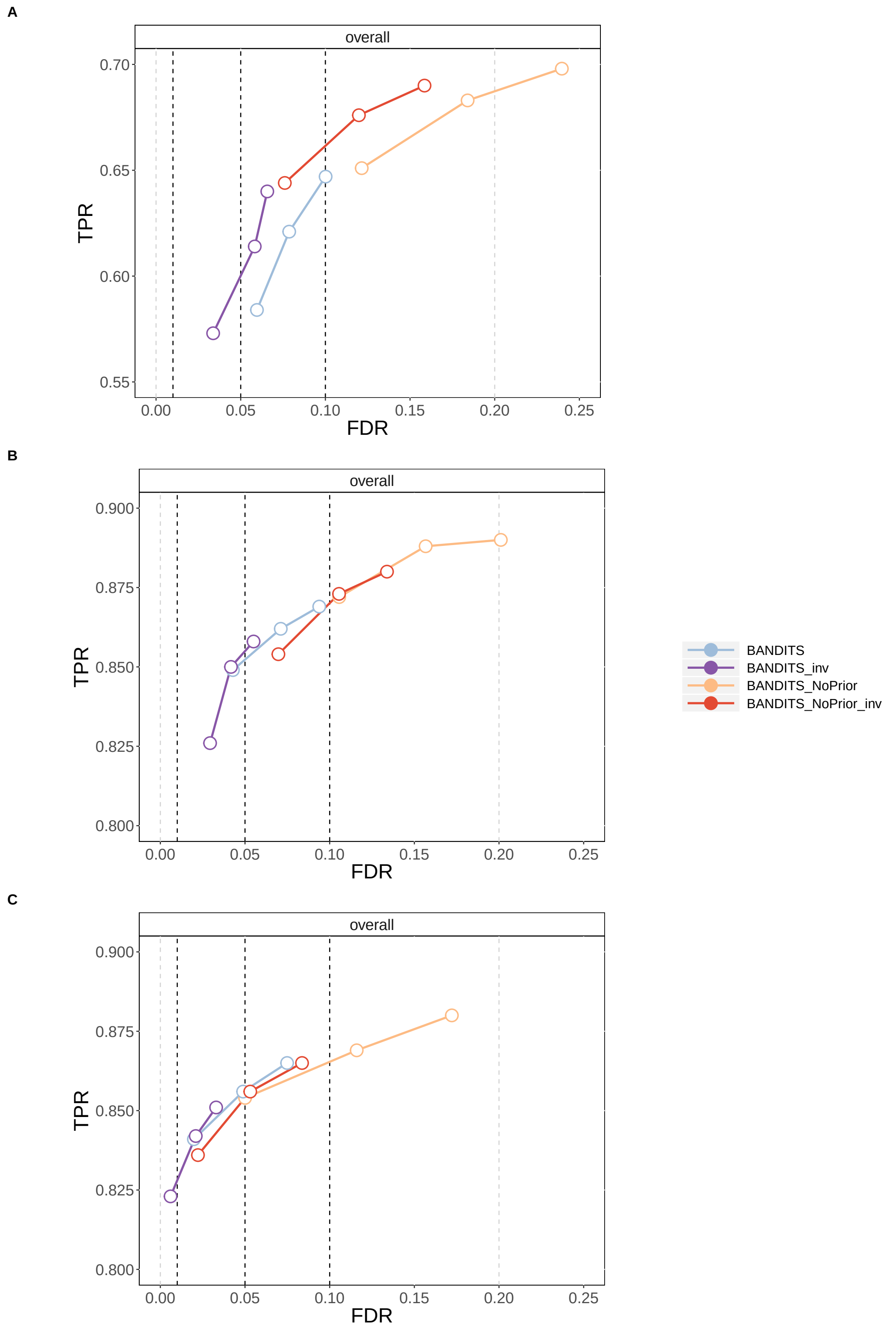

**Figure S19** TPR vs. FDR for gene-level testing for the three 2-group comparison simulation studies. "BANDITS_NoPrior" refers to BANDITS being run with vaguely-informative prior (default when no informative prior is provided). A) 3 vs. 3 simulation study; B) 6 vs. 6 simulation study; C) 6 vs. 6 simulation study with transcript pre-filtering (transcripts with at least 10 counts and an average relative abundance of 0.01). Circles indicate observed FDR for 0.01, 0.05 and 0.1 significance thresholds.

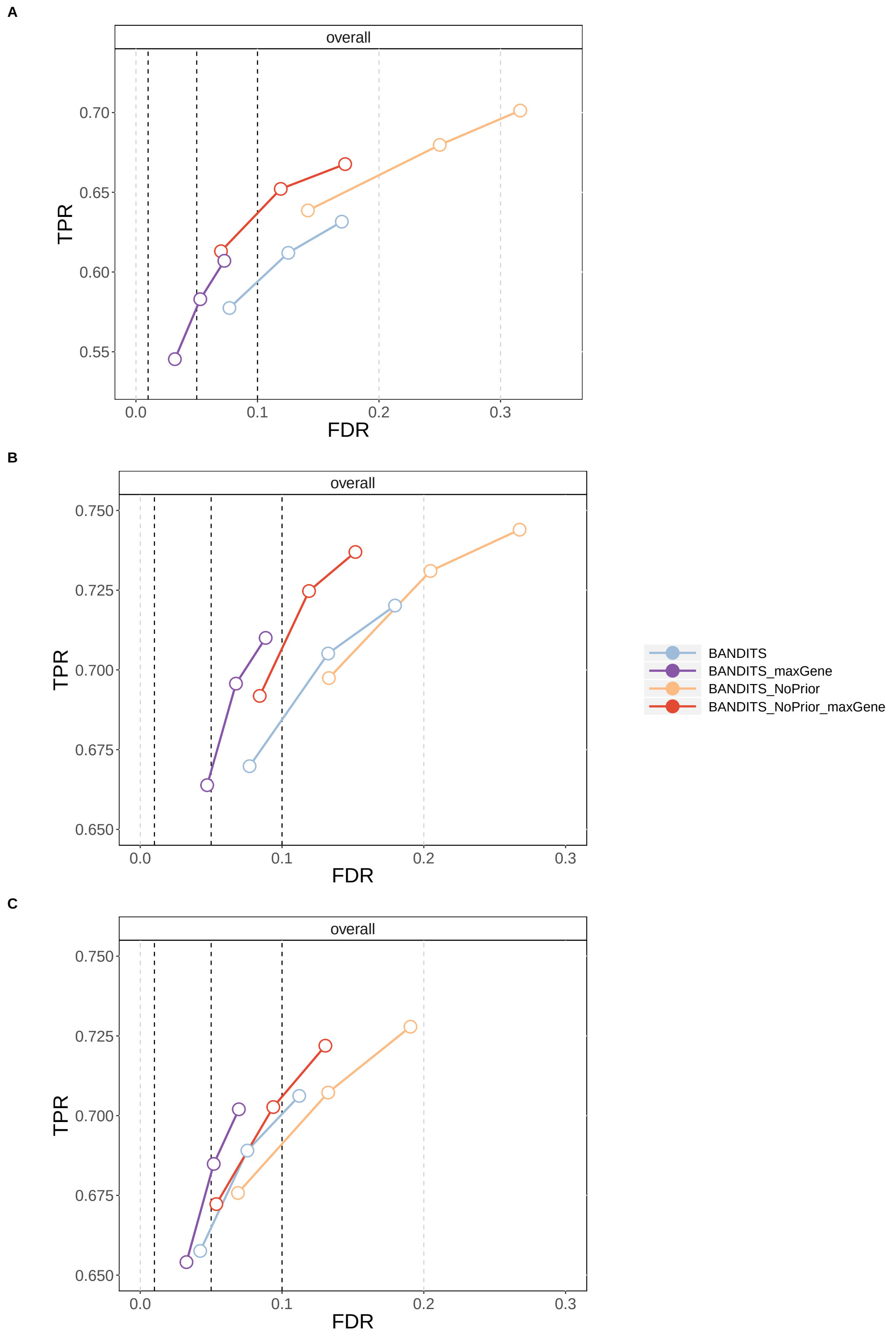

**Figure S20** TPR vs. FDR for transcript-level testing for the three 2-group comparison simulation studies. "BANDITS_NoPrior" refers to BANDITS being run with vaguely-informative prior (default when no informative prior is provided). A) 3 vs. 3 simulation study; B) 6 vs. 6 simulation study; C) 6 vs. 6 simulation study with transcript pre-filtering (transcripts with at least 10 counts and an average relative abundance of 0.01). Circles indicate observed FDR for 0.01, 0.05 and 0.1 significance thresholds.

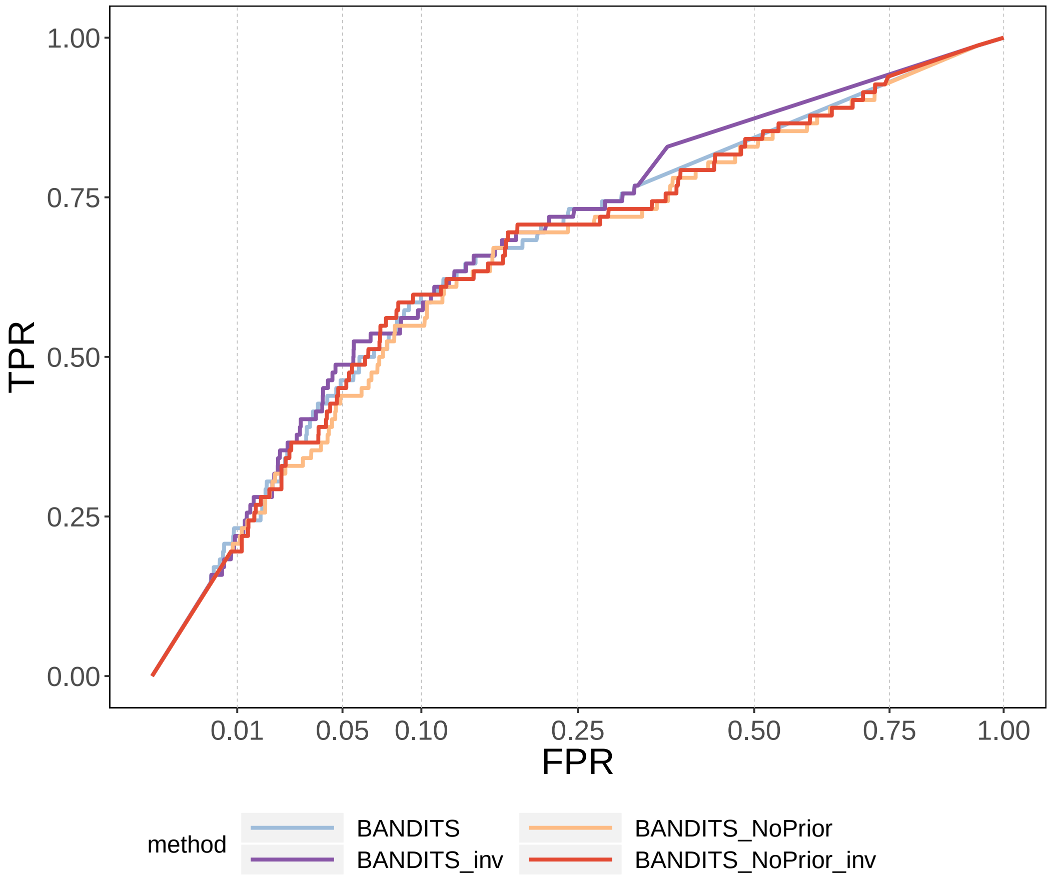

**Figure S21** ROC curve (TPR vs. FPR) for gene-level testing in the "Best et al." experimental dataset. "BANDITS_NoPrior" refers to BANDITS being run with vaguely-informative prior (default when no informative prior is provided).

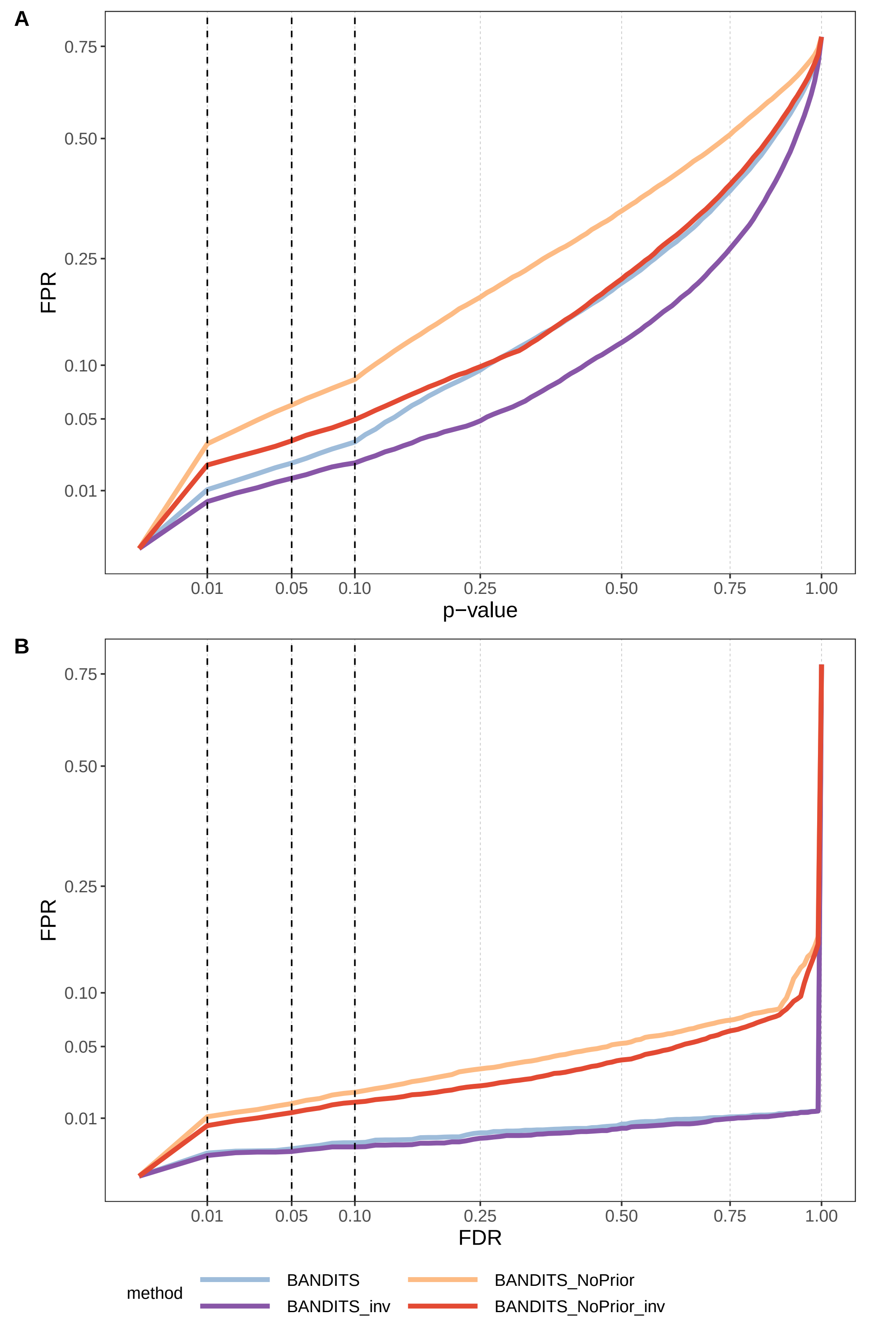

**Figure S22** FPRs for gene-level testing in the null experimental dataset. "BANDITS_NoPrior" refers to BANDITS being run with vaguely-informative prior (default when no informative prior is provided). A) FPR vs. p-value; B) FPR vs. FDR.

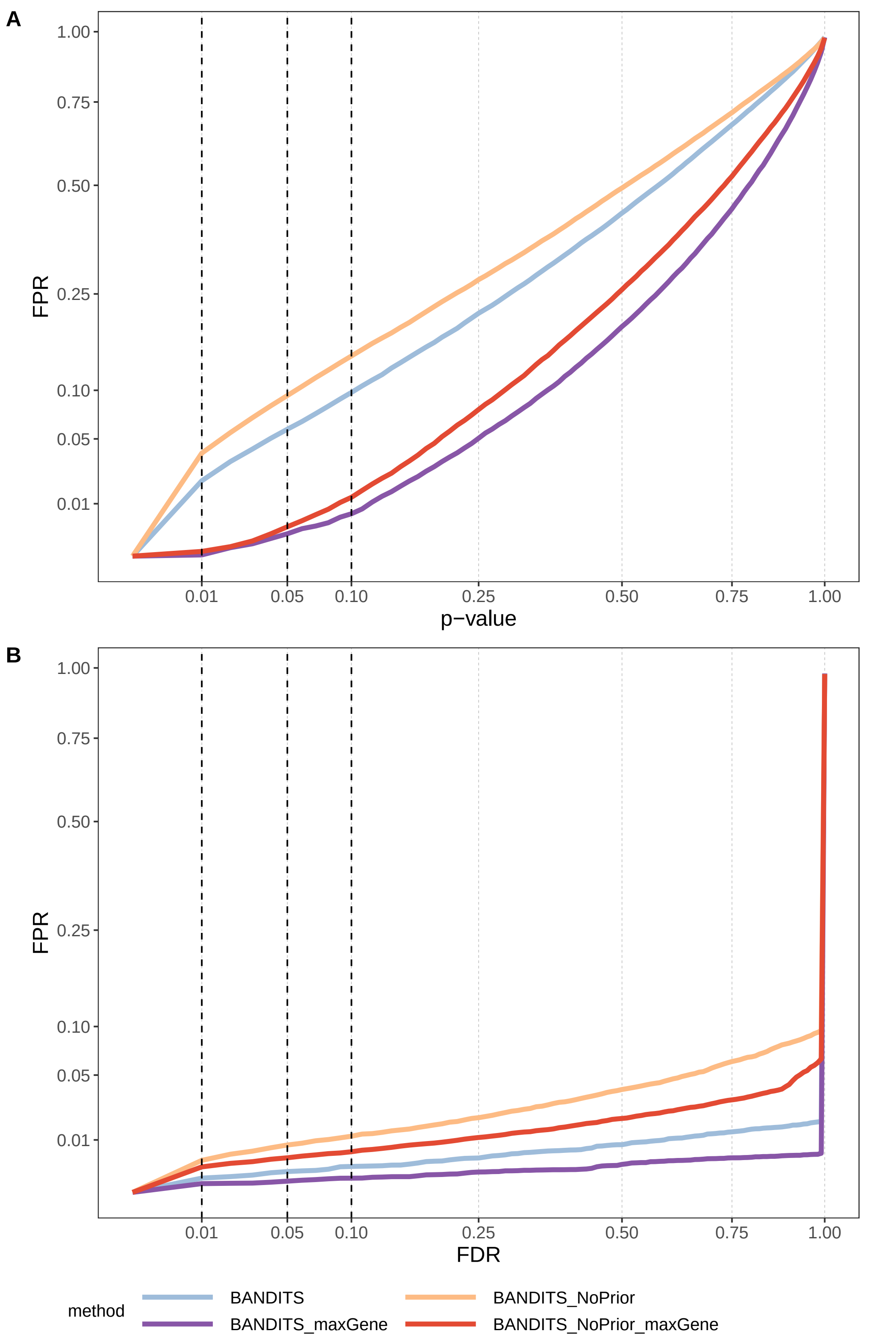

**Figure S23** FPRs for transcript-level testing in the null experimental dataset. "BANDITS_NoPrior" refers to BANDITS being run with vaguely-informative prior (default when no informative prior is provided). A) FPR vs. p-value; B) FPR vs. FDR.

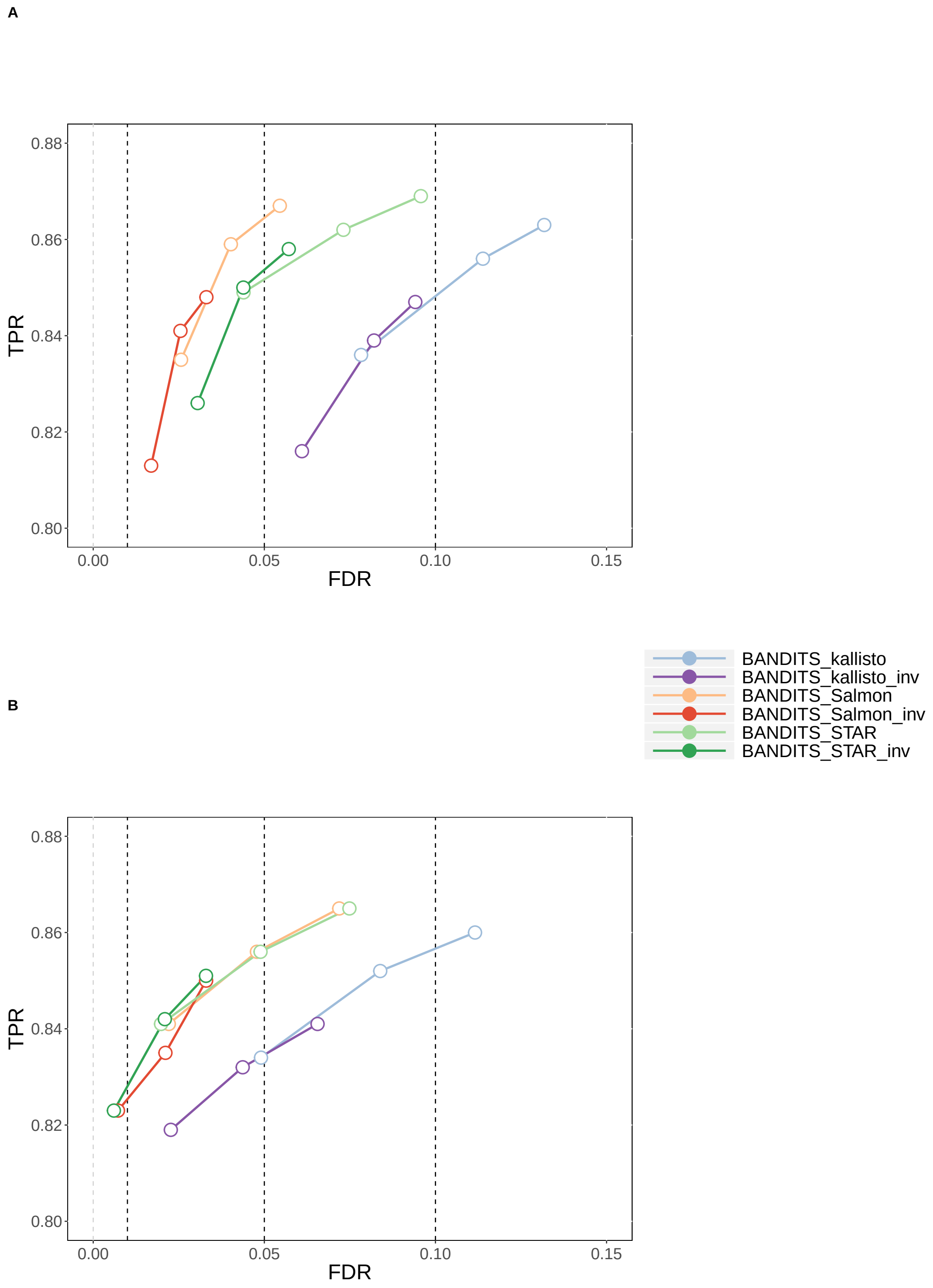

**Figure S24** TPR vs. FDR for gene-level testing for the 2-group comparison simulation study. “kallisto”, “Salmon” and “STAR” refer to the alignment mode. A) 6 vs. 6 simulation study; B) 6 vs. 6 simulation study with transcript pre-filtering (transcripts with at least 10 counts and an average relative abundance of 0.01). Circles indicate observed FDR for 0.01, 0.05 and 0.1 significance thresholds.

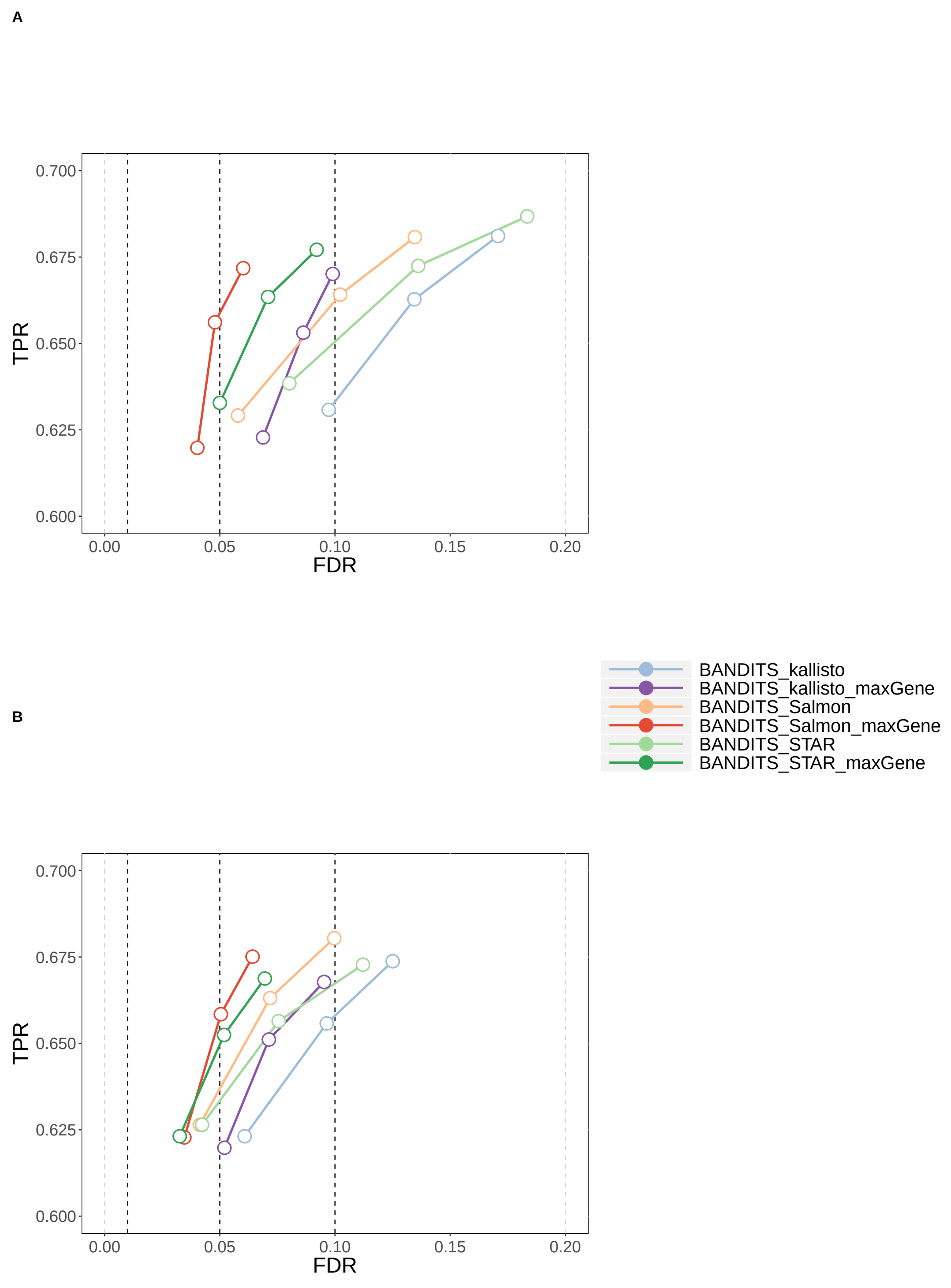

**Figure S25** TPR vs. FDR for transcript-level testing for the 2-group comparison simulation study. "kallisto", "Salmon" and "STAR" refer to the alignment mode. A) 6 vs. 6 simulation study; B) 6 vs. 6 simulation study with transcript pre-filtering (transcripts with at least 10 counts and an average relative abundance of 0.01). Circles indicate observed FDR for 0.01, 0.05 and 0.1 significance thresholds.

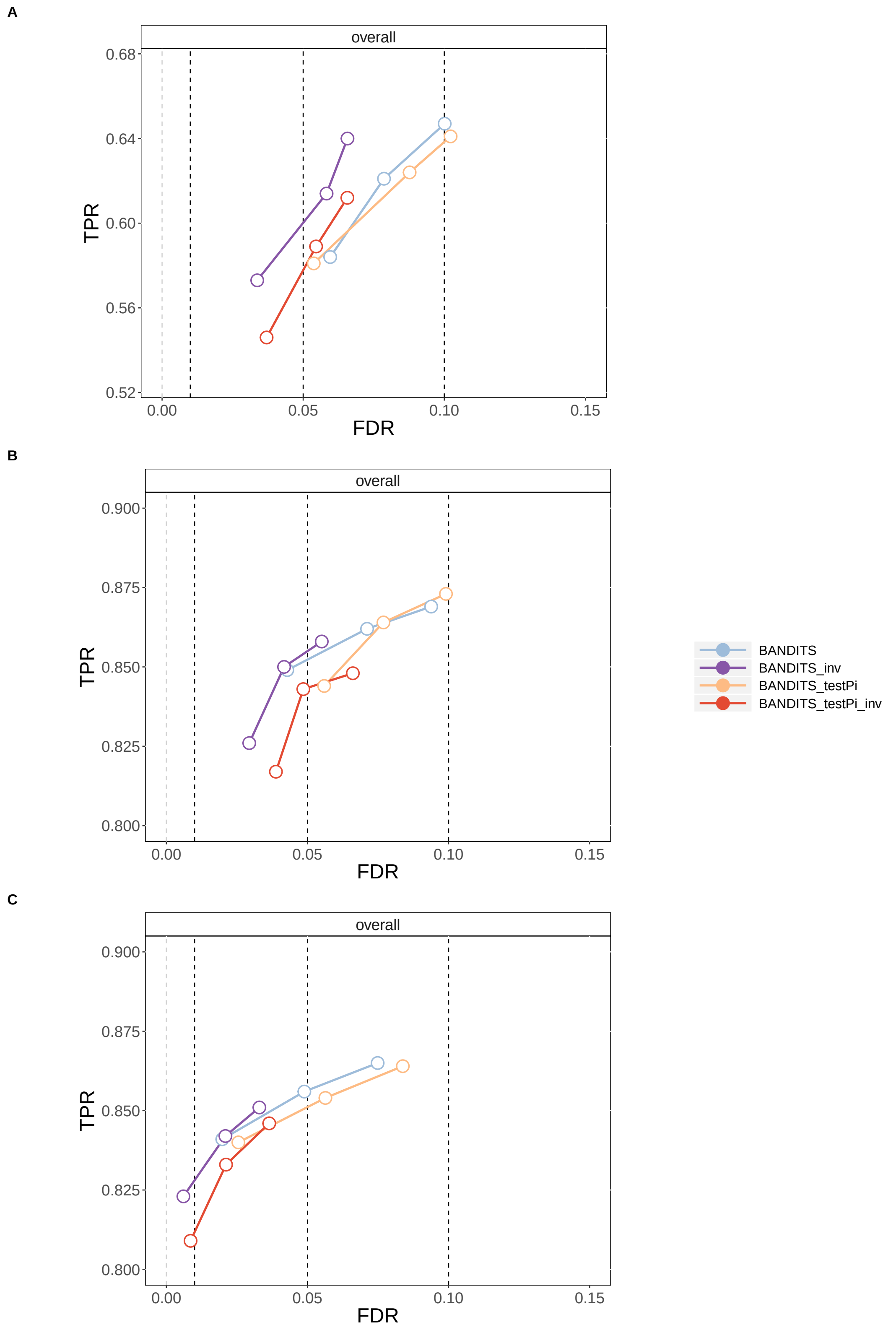

**Figure S26** TPR vs. FDR for gene-level testing for the three 2-group comparison simulation studies. "BANDITS_testPi" refers to a modified version of BANDITS to test the original Dirichlet-multinomial parameter, $\pi$, without normalizing for the transcript effective lengths. A) 3 vs. 3 simulation study; B) 6 vs. 6 simulation study; C) 6 vs. 6 simulation study with transcript pre-filtering (transcripts with at least 10 counts and an average relative abundance of 0.01). Circles indicate observed FDR for 0.01, 0.05 and 0.1 significance thresholds.

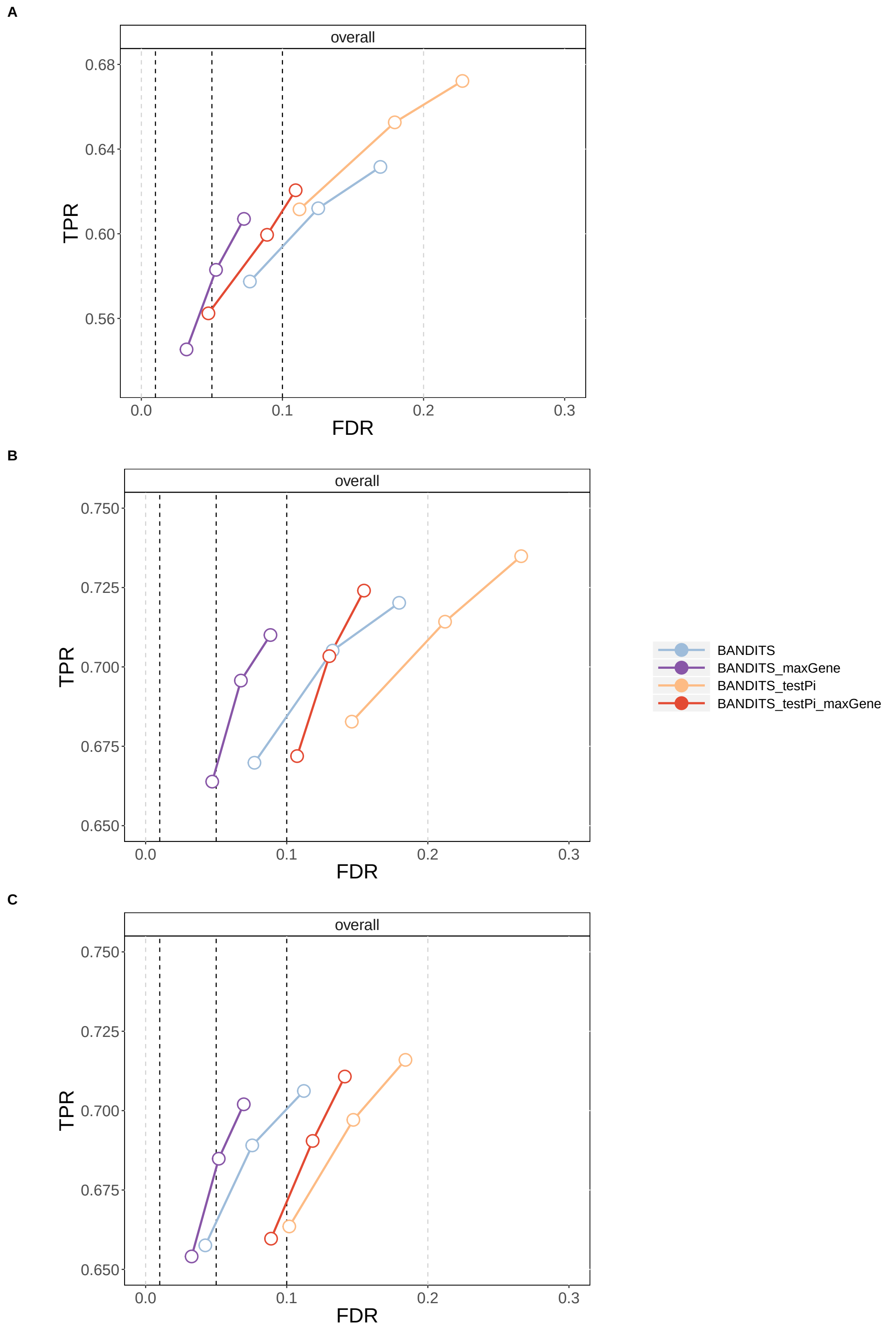

**Figure S27** TPR vs. FDR for transcript-level testing for the three 2-group comparison simulation studies. “BANDITS_testPi” refers to a modified version of BANDITS to test the original Dirichlet-multinomial parameter, $\pi$, without normalizing for the transcript effective lengths. A) 3 vs. 3 simulation study; B) 6 vs. 6 simulation study; C) 6 vs. 6 simulation study with transcript pre-filtering (transcripts with at least 10 counts and an average relative abundance of 0.01). Circles indicate observed FDR for 0.01, 0.05 and 0.1 significance thresholds.

**Author details**

[1]Institute of Molecular Life Sciences, University of Zurich, Winterthurerstrasse 190, 8057, Zurich, Switzerland. [2]SIB Swiss Institute of Bioinformatics, 8057, Zurich, Switzerland.
